# In C57Bl6 Mice, Obesity and Subsequent Weight Loss Negatively Affected the Skeleton and Shifted the Cortical Bone Metabolome

**DOI:** 10.1101/2025.05.09.653133

**Authors:** Carolyn Chlebek, Casey McAndrews, Benjamin Aaronson, Hope D. Welhaven, Kanglun Yu, Samantha N. Costa, Joseph Shaver, Sophia Silvia, Victoria DeMambro, Ronald K. June, Meghan E. McGee-Lawrence, Clifford J. Rosen

## Abstract

Obesity and calorie restriction each negatively affect skeletal health. Despite the negative effects of weight loss on the skeleton, obese patients are advised to lose weight via calorie restriction. Additionally, obesity and weight loss individually alter both whole-body and local metabolism. Little is known about bone quality and changes to the cortical metabolome following calorie restriction in obese preclinical models. We hypothesized that caloric restriction would worsen bone quality in obese mice by shifting the cortical bone metabolome. To induce obesity, 8-week-old male and female C57BL6/J mice received 60% high-fat diet for 12 weeks. From 20 to 30 weeks of age, mice either remained obese or lost weight through 30% caloric restriction. Control animals received a 10% low-fat diet. Bodyweight and fat mass were increased by obesity and decreased with calorie restriction. Similarly, glucose and insulin tolerance were worsened with obesity but improved by weight loss. Compared to obesity, calorie restriction elicited more bone loss in both cortical and trabecular compartments. Weight loss also reduced bone formation. Both obesity and subsequent calorie restriction altered the cortical bone metabolome in a sex-dependent manner. Metabolic pathways altered with diet generally mapped to amino acid or fatty acid metabolism. In males, weight loss was associated with a downregulation of pathways related to tryptophan, tyrosine, ubiquinone, and fatty acids. In females, calorie restriction downregulated taurine and hypotaurine metabolism but upregulated pyrimidine metabolism, nicotinate and nicotinamide metabolism, and pantothenate and CoA biosynthesis. Our findings highlight the negative effects of obesity and subsequent caloric restriction on the skeleton. Despite improvements in components of systemic metabolism, caloric restriction in obese preclinical models did not restore bone morphology or the cortical metabolome to control conditions.

## Introduction

Obesity is associated with a wide range of negative health outcomes, including increased fracture risk^1,2^. The number of patients diagnosed with obesity in the United States is rising^3^, which will lead to more fractures and add to the healthcare burden. In addition to increased fracture risk, patients with obesity are more likely to be diagnosed with advanced osteoarthritis, requiring surgical intervention^4^. Total hip and total knee replacements are common procedures used to improve pain-related symptoms and mobility. These surgeries rely on osseointegration, or the ingrowth of both cortical and trabecular bone onto the implant surface^5^, and thus bone quality may affect surgical outcomes, particularly in patients with obesity. Weight loss is often recommended for patients with obesity and can be achieved via a calorie-restricted diet, exercise, anti-obesity medications, or bariatric surgery. Due to the noninvasive nature of lifestyle interventions, diets such as calorie restriction are often first-line clinical recommendations for weight loss^6^.

Diet-induced obesity in the mouse elicits skeletal changes that mirror clinical findings and thus serve as a useful model of obesity. In high-fat diet-fed mice, trabecular bone morphology is negatively altered, with obesity-induced reductions in bone volume fraction and trabecular number^7–9^. Cortical bone mass also is decreased by obesity, with fewer morphological changes than trabecular bone^7–9^. Additionally, in humans and mice, obesity increases bone marrow adipose tissue accumulation^9–11^, which is negatively correlated with skeletal health^11^.

In the absence of obesity, calorie restriction has been linked to bone loss, both clinically and in rodent models. When calories are restricted in individuals without obesity, bone mineral density is reduced^12,13^. Similarly, in nonobese mice, calorie restriction decreases cortical bone mass and suppresses bone formation^14–18^. The effect of calorie restriction on trabecular bone is less clear; some studies report that trabecular bone negatively affected by caloric restriction^14,15^ whereas other researchers report improved trabecular bone morphology^16–18^. Paradoxically, bone marrow adipose tissue is increased by calorie restriction in nonobese murine models^15,18^ and following weight loss in nonobese humans^19,20^.

Research investigating the effect of weight loss on the skeleton after obesity is limited. In humans with obesity, following weight loss, bone mineral density is increased in males but decreased in females^21^, suggesting sex-dependent effects on bone quality. Similarly, obese mice that were transitioned from a high-to low-fat diet lost weight, but the effects of diet change on bone quality were sex-dependent^9,22^. Following weight loss with a low-fat diet, formerly obese mice retained obesity-induced skeletal deficits^9,22^.

Both obesity and weight loss are associated with shifts in whole-body and local metabolism. Obesity increases circulating glucose levels and negatively affects insulin-stimulated glucose uptake in several organs, including the bone marrow^23^. When obese individuals lose weight, fasting blood glucose is reduced and insulin sensitivity increased^24^. Interestingly, circulating metabolites also shift in obese individuals who alter their diet to achieve weight loss^25^. In addition to whole-body metabolic changes elicited by obesity and subsequent weight loss, cellular metabolism within bone cells is likely also affected. Mitochondrial dysfunction is evident when osteoblasts are differentiated in high glucose media, a method to simulate obesity *in vitro*^26^. Similarly, culturing osteocytes in high fatty acid or high glucose conditions reduces mitochondrial activity and increases cellular reactive oxygen species^27^.

We sought to understand the skeletal changes following caloric restriction in obese preclinical models. We hypothesized that caloric restriction would worsen bone quality in obese mice, independent of sex. Obesity-induced skeletal deficits were confirmed prior to weight loss, and components of whole-body metabolism were evaluated both during obesity and following caloric restriction. In addition to evaluating diet-induced changes to skeletal morphology and bone formation, we also investigated whether obesity and subsequent caloric restriction altered the cortical bone metabolome.

## Materials and Methods

### Mice

Male and female C57BL/6J mice aged 5 weeks (Jackson Laboratories, Bar Harbor, ME) were acclimated for three weeks prior to the experiment. All mouse procedures were performed under the approval of the Institutional Animal Care and Use Committee (IACUC) at the MaineHealth Institute for Research. Throughout the study, mice were housed in 14-hour light:10-hour dark cycles. Outside of fasting periods, all animals had ad libitum access to food and water. Bodyweight was recorded at baseline and weekly thereafter.

### Dual energy x-ray absorptiometry

Body mass, composition, bone mineral content (BMC), and bone mineral density (BMD) were measured at baseline and at the end of each diet phase using dual energy x-ray absorptiometry (Faxitron UltraFocus-DXA, Hologic, Marlborough, MA). Calibration was performed using phantom standards provided by the manufacturer. The mouse cranium was removed from analysis. Areal bone mineral content, bone mineral density, fat mass, percent fat, and lean mass were calculated.

### Obesity Induction

During the High Fat Feeding Phase, mice were housed in groups of two or three per cage. At 8 weeks of age, the High Fat Feeding Phase was initiated. Cages were randomly assigned to receive a 60% high fat diet, containing 60% of calories from fat (n=34) or a 10% kcal low fat diet, containing 10% of calories from fat (n=18) (D12492i and D12450Ji, respectively; Research Diets Inc, New Brunswick, NJ). Male mice received at least 3.5 g of diet per mouse per day; females had at least 3 g per mouse per day. Low fat diet consumption was evaluated weekly. High fat diet was completely replaced every 2-3 days.

Following 12 weeks of high- or low-fat feeding, mice were determined to be obese or nonobese by evaluating bodyweight, fat mass, and percent body fat (Figure S1). Based on these criteria, seven female mice were excluded at the end of the High Fat Feeding Phase due to failure to become obese. To confirm the skeletal phenotype in mice at the end of the High Fat Feeding Phase, 5 mice per group were euthanized in both males and females. One male mouse was euthanized prior to the conclusion of the High Fat Feeding Phase due to a bad reaction to an insulin injection.

### Weight Loss Induction

Following obesity induction, obese mice were randomly assigned to remain on the high fat diet (HFD) or were assigned to the diet-induced weight loss (HFD-CR). At the conclusion of the High Fat Feeding Phase, all mice were single housed. HFD animals continued to receive 60% high fat diet. HFD-CR mice received the low fat diet for 2 weeks (Stabilization Phase) and were then transitioned to a 30% calorie-restricted diet for 8 weeks (Caloric Restriction Phase) (Figure 1A). All mice were euthanized at the end of the Caloric Restriction Phase, at 30 weeks of age.

**Figure 1.**
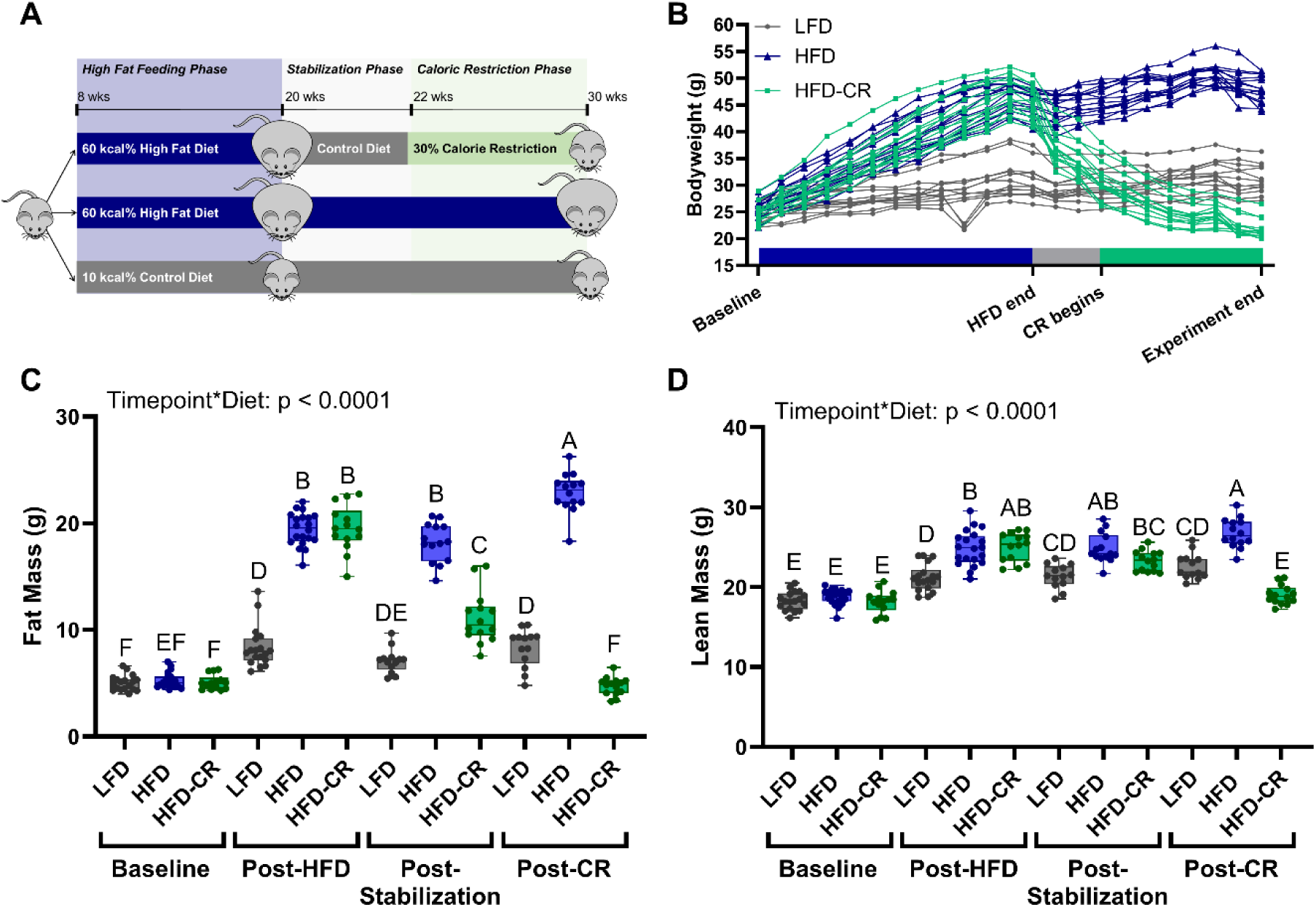
Body mass increased during high fat feeding and decreased during caloric restriction. (A) Obesity was induced in 8-week-old male and female mice with a diet containing 60% kcal from fat (HFD) during the High Fat Feeding Phase. After 12 weeks of high fat feeding, mice meeting the obesity criteria were randomly assigned to lose weight with caloric restriction or remain obese. To induce weight loss, animals were transitioned to a diet containing 10% kcal from fat (LFD) for 2 weeks, followed by 8 weeks of caloric restriction. A final control group received LFD for the duration of the experiment. (B) During the High Fat Feeding Phase, males receiving high fat diet gained bodyweight. During the Caloric Restriction Phase, calorie restricted mice lost weight. (C) Body scans were obtained at baseline and at the end of each diet phase using DXA. HFD increased fat mass, but stabilization reduced fat mass. Subsequent caloric restriction further decreased fat mass. (D) Lean mass was increased by HFD but decreased following caloric restriction. Fat and lean mass were analyzed by two-way ANOVA with factors of timepoint and diet. Significant interaction terms are displayed. Tukey post-hoc analysis determined differences between groups. p≤0.05 for all effects displayed. Letters denote statistical differences: A>B>C>D>E>F.

When fed at 30% restriction, the calorie restriction diet (D15032801i; Research Diets Inc, New Brunswick, NJ) contains equal amounts of vitamins and minerals to that consumed by control mice. Mice receiving a low fat diet during obesity induction remained on the same diet for the study duration (Control). Weekly food consumption in the control group was used to determine the amount of calorie restriction food for HFD-CR mice in the subsequent week. HFD-CR mice were fed within the same two-hour window each day. Two female HFD-CR mice were euthanized prior to the end of the study due to lack of food consumption. An additional two female mice were euthanized due to dermatitis.

### Glucose and Insulin Tolerance Testing

Glucose and insulin tolerance were assessed at the end of the High Fat Feeding and Caloric Restriction Phases. Mice were fasted for 6 hours with free access to water. Fasting blood glucose was then recorded (AlphaTrak 2, Zoetis, Parsippany-Troy Hills, NJ). Mice were then injected intraperitoneally with glucose (1g/kg bodyweight) or insulin (1 U/kg bodyweight). Blood glucose was recorded at 15, 30, 45, 60 and 120 minutes following injection. For glucose tolerance tests, blood glucose was plotted over time. For insulin tolerance tests, blood glucose was first normalized to the fasting blood glucose and then plotted over time. Area under the curve was calculated.

### Indirect Calorimetry

One week prior to euthanasia, 8 animals from each group were housed in metabolic cages (Promethion cages, Sable Systems Intl., North Las Vegas, NV). Metabolic cages contained standard bedding, a food hopper, water bottle, a house-like enrichment tube for body mass measurements, and a 11.5 cm running wheel that recorded revolutions. Following a 24 hour acclimation in the metabolic cages, data was collected for 72 hours. Data was analyzed separately for day, night, and 24-hour cycles. Promethion Live software (v23.0.5) was used for data acquisition and instrument control. Raw data was processed with Macro Interpreter (v22.10). Respiratory quotient and energy expenditure were determined by assessment of respiratory gases, using standard equations^17,28^ (Promethion Core™ CCF, Sable Systems Intl.). Ambulatory activity and the position of each mouse was determined with XYZ beam arrays (beam spacing of 0.25 cm).

### Adipose Depot Evaluation

At euthanasia, both inguinal fat pads and intrascapular brown adipose depots were collected and weighed. Tissues were then fixed in 10% formalin for 48 hours on an orbital shaker. Following fixation, adipose depots were transferred to 70% ethanol. Inguinal fat pads from five mice per group were then sectioned and stained for hematoxylin and eosin. adiposity was quantified in ImageJ (version 2.14.0/1.54f) using a region of interest measuring 263 µm^2^. Adipocyte number and size were averaged across two sequential sections per animal^29^.

### Dynamic Histomorphometry

Prior to euthanasia (−10 days, -3 days), mice received calcein injections (10 mg/kg). One tibia from each mouse was embedded in plastic and sectioned for dynamic histomorphometry. Cortical bone was sectioned in the transverse plane whereas trabecular bone was sectioned in the coronal plane. A single user calculated bone surface, single labels, and double labels in a blinded fashion for cortical bone. A second user quantified bone surface, single labels, and double labels in trabecular bone. Measurements of mineral apposition rate (MAR), mineralized surface per bone surface (MS/BS), and bone formation rate (BFR/BS) were calculated for trabecular and cortical bone BioQuant (BIOQUANT OSTEO 2021, Nashville, TN). Cortical endosteal and periosteal surfaces were evaluated separately. For trabecular bone samples, the region of interest consisted of a 3×4 grid of 800×500 µm panels starting at the bottom of the growth plate, excluding anything within 200 µm of the cortical bone.

### Micro-computed tomography

At euthanasia, one tibia per mouse was collected for micro-computed tomography (µCT). Bones were fixed in 10% formalin and placed on a rocker for 24 hours. Bones were then rinsed and transferred to 70% ethanol, then stored at 4°C. In accordance with the American Society for Bone and Mineral Research guidelines^30^, all bones were scanned using the same instrument (vivaCT 40, Scanco Medical AG, Brüttisellen, Switzerland) under the same conditions. Scans were acquired using a 10.5 μm^3^ isotropic voxel size, 70 kVp peak x-ray tube intensity, 114 mA x-ray tube current, and 250 ms integration time. All scans were subjected to Gaussian filtration and segmentation. All analyses were performed using the Scanco software.

### Tibiae Histology and Immunohistochemistry

Following analysis with µCT, five bones per group were decalcified in 14% EDTA for two weeks. Tibiae were then paraffin-embedded and sectioned sagittally at 5 µm thickness. Sections were stained for hematoxylin and eosin, TRAP (Sigma-Adrich 387A), or anti pro-collagen-1 specific antibodies (Developmental Studies Hybridoma Bank SP1.D8). In hematoxylin and eosin-stained sections, bone marrow adiposity was quantified in ImageJ using a region of interest beginning 200 um below the growth plate and extending 3000um into the marrow cavity. Using previously established methods^29^, adipocyte number and size were averaged across two sequential sections per animal.

To evaluate osteoblast and osteoclast number, metaphyseal endocortical bone surface and trabecular bone were analyzed separately. Bone-lining, cuboidal cells stained by procollagen-1 were quantified and normalized to bone surface. Similarly, the number of TRAP-positive cells was normalized to the bone surface and quantified.

### Serum Collection and Analysis

Following euthanasia, blood was collected and stored at 8° C for 24 hours. Samples were then centrifuged and serum collected and stored at -80°C. Serum CTx and P1NP were measured by ELISA according to manufacturer’s instructions (IDS AC-06F1 and IDS AC-33f1, Bolden Colliery, United Kingdom).

### Osteoclast Culture

At euthanasia, five mice were used for osteoclast cell culture. Animals were not pooled for cell culture. Bone marrow was removed from femur and pelvic bones via centrifugation. Cells were spun directly into tissue culture media (αMEM, 10% fetal bovine serum and 1% penicillin/streptomycin). Pelleted marrow was physically separated, and cells were then incubated with RT ACK lysing buffer for a maximum of three minutes. Cells were strained and pelleted, then resuspended in culture media and plated in T75 flasks. After 48 hours, non-adherent cells were collected and plated in 48-well plates at 1 million cells/well. Three wells per mouse were plated and cultured in osteoclast differentiation media (tissue culture media with 50 ng/mL RANKL and 25 ng/mL M-CSF). Following 7 days in culture, cells were fixed and stained for TRAP. TRAP-positive cells with three or more nuclei were defined as osteoclasts; osteoclast size and number were quantified in ImageJ.

### Metabolomics

At euthanasia, bone marrow was removed via centrifugation from femurs in all animals. Trabecular bone was removed from the femur and the cortical shell flash frozen in liquid nitrogen. A subset of femurs collected at the end of the Caloric Restriction Phase were included for metabolomic analysis (Figure S2). Using a bead mill, bone was pulverized in a 70:30 methanol:acetone solution. Samples were centrifuged and supernatant containing metabolites was extracted. Supernatant was then dried in vacuum concentrator. Metabolites were then resuspended in 1:1 acetonitrile:water. All solvents used were high-performance liquid chromatography grade.

Following metabolite extraction^31^, samples were analyzed by untargeted liquid chromatography-mass spectrometry (LC-MS) using Waters I-Class Ultra-High Performance Liquid Chromatography coupled to a Waters Synapt-XS Q-IMS-TOF in positive mode (Waters, Milford, MA). A Cogent Diamond Hydride hydrophilic interaction liquid chromatography (HILIC) chromatography column was utilized (2.2μM, 120 Å, 150 mm X 2.1 mm; MicroSolv, Leland, NC, USA). 5 μL of sample was injected and blank samples were analyzed every 10 samples for quality control to prevent spectral drift and contamination. LC-MS data was then exported and converted for analysis using Progenesis.

Metabolomic data was assessed with MetaboAnalyst^31^. Normalized abundances were log transformed, standardized and auto-scaled. For all metabolic analysis, male and female mice were analyzed separately. To determine whether largescale shifts in metabotype occurred following weight gain and weight loss, partial least squares-discriminant analysis was performed. To identify metabolic pathways altered by diet, HFD and HFD-CR metabolite intensities were normalized to the median intensity in LFD animals and hierarchical clustering performed. The metabolites comprising each cluster were then correlated to metabolic pathways using MetaboAnalyst’s MS Peaks to Pathways feature with the *mummichog* algorithm.

### Statistical Analyses

For all measures, male and female mice were analyzed separately. Longitudinal data was analyzed using a two-way ANOVA with factors of diet and timepoint (bone mineral density, body composition). Tukey’s post-hoc tests determined significant differences between groups. Glucose and insulin tolerance tests performed at the end of each diet phase were analyzed separately to avoid any confounding factors, such as differences in experimental handlers or room conditions. One-way ANOVA for diet was used to assess differences for area under the curve for glucose and insulin tolerance tests, as well as ambulatory data collected during metabolic cage housing. For energy expenditure data collected with indirect calorimetry, diet differences were determined with ANCOVA, to correct for differences lean mass.

To determine differences following obesity induction, t-tests were used in for the small cohort of euthanized animals (peripheral adipose depots, bone morphology, dynamic histomorphometry, osteoblast number, osteoclast number, marrow adiposity).

To determine differences following weight loss, data were analyzed using a one-way ANOVA for diet (bone morphology, dynamic histomorphometry, P1NP, osteoblast number, CTx, osteoclast number, osteoclastogenesis, marrow adiposity). Tukey’s post-hoc tests determined significant differences between groups.

In all tests, significance was set at p<0.05. All statistical analyses were performed in R.

## Results

### High fat diet and subsequent caloric restriction changed body composition and whole-body metabolism

Whole-body metabolism worsened with high fat feeding but improved with caloric restriction in both male and female mice. Compared to LFD control animals, HFD and HFD-CR mice gained body weight and fat mass during high fat feeding (Figure 1B). Caloric restriction reduced body weight and fat mass in HFD-CR compared to both HFD and LFD mice (Figure 1B-C, S3A-B). Lean mass increased during high fat feeding and decreased with caloric restriction (Figure 1D, S3C).

Diet altered both glucose and insulin tolerance. In both sexes, fasting blood glucose was elevated by high fat feeding but reduced by caloric restriction (Figure S4). Compared to control mice, glucose tolerance was worsened in HFD and HFD-CR mice after the High Fat Feeding Phase in both male and female mice (Figure S5A). In both sexes, caloric restriction improved glucose tolerance in HFD-CR compared to both LFD and HFD animals (Figure S5B). After high fat feeding, insulin tolerance was worsened in male mice, but not altered in females (Figure S5C). By the end of the study, HFD females had worse insulin tolerance compared to LFD mice (Figure S5D). Following caloric restriction, HFD-CR mice had improved insulin tolerance compared to both HFD and LFD mice, in both sexes (Figure S5D).

Whole-body metabolism was further analyzed with indirect calorimetry. In male mice, after accounting for lean mass, energy expenditure did not vary between groups. In females, 24-hour energy expenditure was significantly higher in HFD compared to LFD mice, even after accounting for differences in lean mass. In both sexes, ambulatory patterns varied by diet. Calorie-restricted mice ran further and faster compared to both HFD and LFD animals (Figure S6A). Compared to HFD animals, HFD-CR mice also walked further and faster within the cage (Figure S6B). Walking distance did not vary between LFD and HFD-CR mice.

### Diet altered the mass and adiposity of white inguinal and bone marrow adipose tissue

Adipose depots were evaluated in a small cohort of mice euthanized after the High Fat Feeding Phase and at the end of the Caloric Restriction Phase (Figure 1A). Diet regimen altered the mass of white adipose depots more than intrascapular brown fat. In both male and female mice, inguinal fat pad weights increased with high fat feeding and decreased with caloric restriction (Table S1, Figure S7). Conversely, after 12 weeks of high fat feeding, brown adipose weight was not changed by HFD in male or female mice (Table S1). In both sexes, after the Caloric Restriction Phase, calorie restricted animals had smaller brown fat pads compared to HFD mice; however, following normalization of brown fat mass to bodyweight, there were no differences between HFD and HFD-CR (Figure S7B-C). In female mice, the proportion of brown adipose mass to bodyweight was reduced by HFD compared to LFD (Figure S7C).

Obesity changed the adiposity of inguinal adipose depots. At the conclusion of the High Fat Feeding Phase, inguinal adipocytes were larger in obese animals compared to controls in both male and female mice (Table S1). Increased adipocyte size was maintained in HFD mice, but HFD-CR animals had smaller adipocytes (Figure S8). This increase in adipocyte size resulted in fewer adipocytes per area (Table S1, Figure S8). In male mice, the percent area occupied by inguinal adipocytes was not altered by 12 weeks of high fat diet; conversely, obese females had a higher percent area occupied by adipocytes compared to LFD controls (Table S1). After the Caloric Restriction Phase, HFD mice had a higher percent area occupied by adipocytes compared to both LFD and HFD-CR (Figure S8).

Adiposity of bone marrow adipose tissue was altered by diet regimen. Following the High Fat Feeding Phase, obesity increased the size of bone marrow adipocytes, but adiposity was not significantly increased (Table S1). Similarly, after the Caloric Restriction Phase, adipocytes in HFD mice were larger than LFD in both sexes (Figure S9). Female HFD-CR mice also had larger adipocytes compared to LFD controls. In male mice, caloric restriction following HFD significantly increased adipocyte number compared to LFD. The percent area occupied by adipocytes was similarly increased in HFD-CR females versus LFD.

### High fat feeding incurred negative skeletal effects

After 12 weeks of high fat feeding, a small subset of mice was euthanized to confirm the skeletal changes induced by obesity. Compared to LFD mice, high fat feeding reduced cortical and trabecular thickness in males (Table S2). Obesity also reduced trabecular bone formation rate and mineralizing surface in male mice, though not statistically different in our small sample size (p < 0.1). In female mice, obese animals had more cortical area compared to LFD. Compared to LFD mice, HFD females had reduced periosteal bone formation and mineral apposition rates, though not statistically different in our small sample size (p < 0.1). In this small cohort, the number of TRAP-or Pro-collagen-1-positive cells on the trabecular and cortical surfaces did not differ between LFD and HFD mice after 12 weeks of high fat diet in either sex.

### Calorie restriction negatively affected skeletal morphology in previously obese mice

Changes in areal bone mineral density (aBMD) and bone mineral content (aBMC) were monitored at the end of each diet phase (Figure S10). Both aBMD and aBMC increased in LFD mice during the High Fat Feeding Phase. Compared to baseline, male mice receiving HFD during the High Fat Feeding Phase had increased aBMC but not aBMD. Conversely, in females, aBMD was not different by diet regiment, but HFD mice had greater aBMC compared to LFD controls. Following the 2-week Stabilization Phase, HFD-CR males had increased aBMD compared to their obese counterparts. Following the Caloric Restriction Phase, compared to HFD males, HFD-CR mice retained increased aBMD. In both males and females, caloric restriction reduced aBMC compared to HFD mice.

Calorie restriction negatively affected skeletal morphology more than obesity alone (Figure S11). At the end of the Caloric Restriction Phase, cortical area, thickness, and area fraction were reduced in HFD-CR mice compared to controls in both sexes (Figure 2). Compared to obese mice, calorie restriction increased marrow area and decreased the moment of inertia in females, but not males. Cortical tissue mineral density was not different between diet groups. In both sexes, caloric restriction reduced tibia length compared to obese mice. In females, total cortical area increased in HFD versus LFD mice, but did not change with HFD-CR. Total cortical area did not differ between diets in male mice.

**Figure 2.**
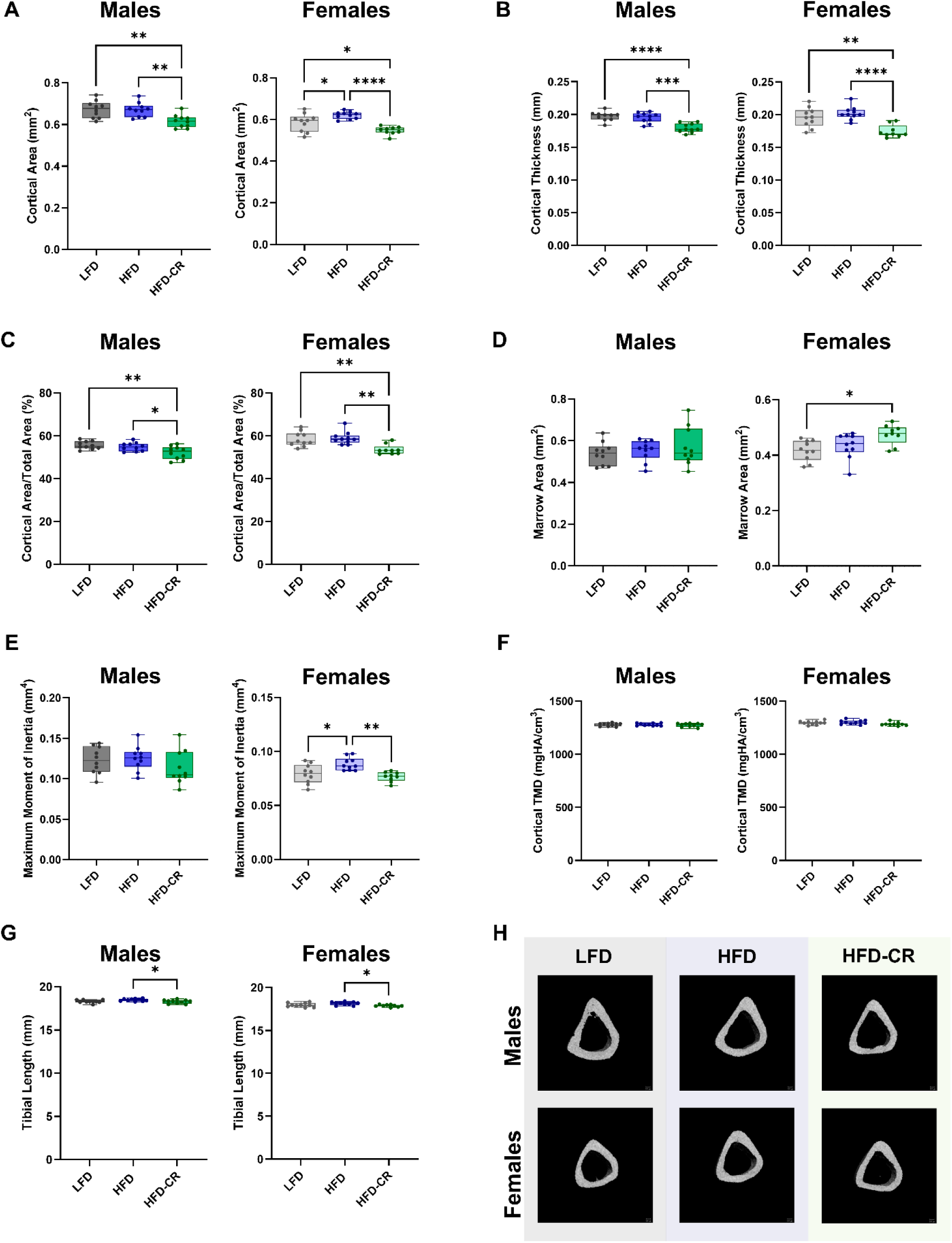
Caloric restriction reduced cortical bone mass in previously obese mice. µCT was used to evaluate cortical bone morphology at the mid-diaphysis. (A) In both male and female mice, calorie restriction reduced cortical area compared to HFD. (B-C) Calorie restriction reduced both cortical thickness and cortical area fraction in both sexes. (D) In females, calorie restriction increased marrow area. Diet did not significantly alter marrow area in males. (E) Compared to their HFD counterparts, calorie restricted females had reduced maximum moment of inertia. Moment of inertia was not different by diet in males. (F) Diet did not change cortical tissue mineral density. (G) Calorie restricted mice had shorter tibiae compared to HFD animals. (H) Representative µCT images. Cortical bone morphology was evaluated by one-way ANOVA with factor of diet. Tukey post-hoc analysis determined differences between groups. p≤0.05 for all effects displayed. * indicates p≤0.05. ** indicates p≤0.01. *** indicates p≤0.001. **** indicates p≤0.0001.

Trabecular bone was altered by HFD in both sexes (Figure 3). In males, bone volume fraction was reduced in both HFD and HFD-CR mice compared to controls. Diet did not change trabecular bone volume fraction in females. Compared to HFD and LFD animals, HFD-CR mice had reduced trabecular thickness in both sexes. Compared to LFD, male HFD and HFD-CR mice had increased trabecular separation but reduced trabecular number. In females, caloric restriction reduced trabecular separation and increased trabecular number in HFD-CR compared to HFD mice. Compared to LFD mice, HFD and HFD-CR reduced trabecular tissue mineral density in male mice. Diet did not change trabecular tissue mineral density in females.

**Figure 3.**
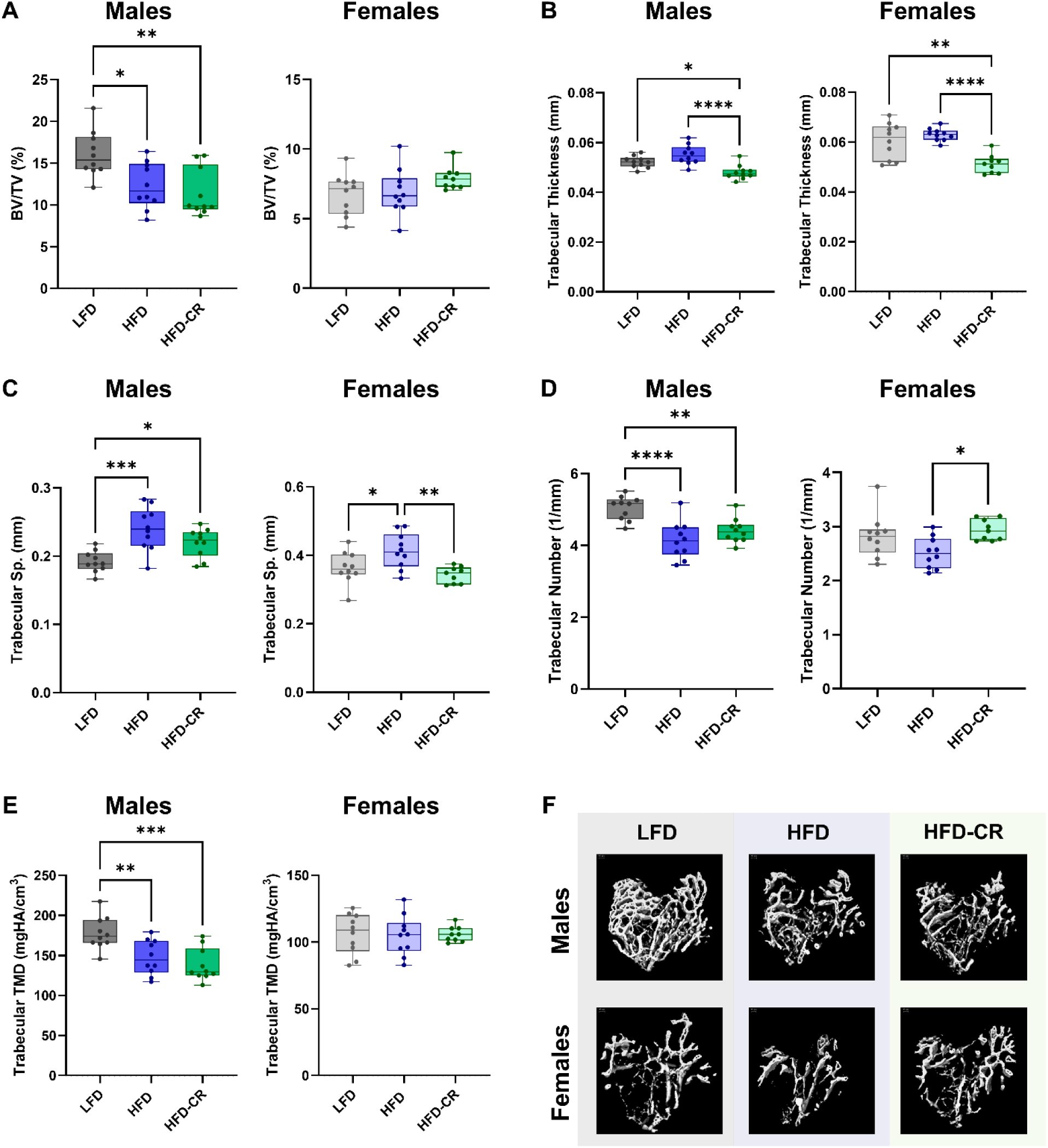
Obesity and subsequent caloric restriction negatively affected trabecular bone mass. µCT was used to evaluate trabecular bone morphology. (A) In male mice, bone volume fraction was reduced in both HFD and HFD-CR animals. Bone volume fraction was not different by diet in female mice. (B) Compared to HFD, calorie restriction reduced trabecular thickness in both sexes. (C) HFD increased trabecular separation in both male and female mice. In females, subsequent caloric restriction increased trabecular spacing compared to HFD mice. (D) In male mice, both HFD and HFD-CR reduced trabecular number. Compared to HFD, HFD-CR increased trabecular number in females. (E) Trabecular tissue mineral density was reduced in both HFD and HFD-CR males but not different in females. (F) Representative µCT images. Trabecular bone morphology was evaluated by one-way ANOVA with factor of diet. Tukey post-hoc analysis determined differences between groups. p≤0.05 for all effects displayed. * indicates p≤0.05. ** indicates p≤0.01. *** indicates p≤0.001. **** indicates p≤0.0001.

### Calorie restriction reduced bone remodeling in both male and female mice

Caloric restriction reduced bone remodeling activities in both the cortical and trabecular compartments. In both sexes, calorie restriction decreased bone formation rates at the periosteal and endosteal surfaces compared to HFD and LFD mice (Figure 4). At the periosteal surface, mineralizing surface and mineral apposition rate were lower in HFD-CR mice compared to HFD in both sexes. In males, endocortical mineralizing surface was reduced in HFD-CR versus LFD. HFD-CR females had decreased endocortical mineralizing surface compared to their obese counterparts. Caloric restriction reduced endocortical mineral apposition rate in male HFD-CR versus HFD. Both HFD and HFD-CR decreased endocortical mineral apposition rate in females.

**Figure 4.**
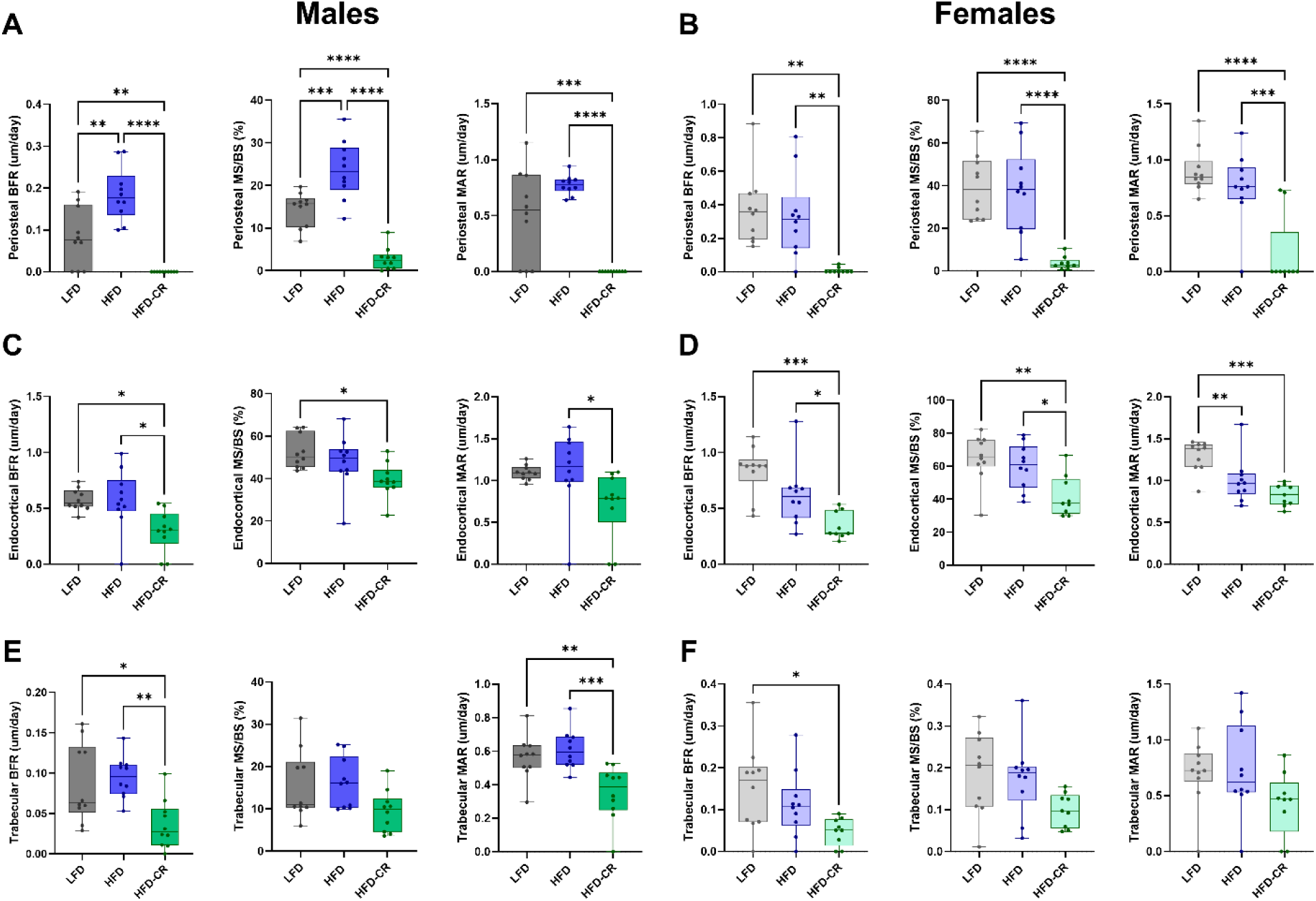
Calorie restriction after obesity reduced bone formation rates. Dynamic histomorphometry was used to evaluate bone formation in cortical and trabecular bone. (A-B) Compared to HFD, calorie restriction reduced bone formation rate, mineralizing surface, and mineral apposition rate at the periosteal surface in both male and female mice. (C-D) At the endosteal surface, bone formation rate was lower in HFD-CR compared to HFD mice in both sexes. In males, endosteal mineralizing surface was lower in HFD-CR versus LFD mice. Compared to HFD, calorie restriction reduced mineral apposition rate males. In females, calorie restriction reduced endosteal mineralizing surface compared to both LFD and HFD mice. Both HFD and HFD-CR females had lower endosteal mineral apposition rates compared to LFD. (E) In males, caloric restriction reduced trabecular bone formation rate and mineral apposition rate compared to both HFD and LFD. Trabecular mineralizing surface was not different across diets in male mice. (F) Compared to LFD, HFD-CR reduced trabecular bone formation rate in female mice. Diet did not change trabecular mineralizing surface or mineral apposition rates in females. Bone formation rate, mineralizing surface, and mineral apposition rate were evaluated by one-way ANOVA with factor of diet. Tukey post-hoc analysis determined differences between groups. p≤0.05 for all effects displayed. * indicates p≤0.05. ** indicates p≤0.01. *** indicates p≤0.001. **** indicates p≤0.0001.

In male mice, trabecular bone formation rate was reduced in HFD-CR compared to HFD and LFD mice. Trabecular bone formation rates were lower in HFD-CR versus LFD females. In both sexes, trabecular mineralizing surface was not altered by diet. Compared to LFD and HFD mice, calorie restriction reduced trabecular mineral apposition rate in males. Trabecular mineral apposition rate did not differ by diet in female mice.

Serum P1NP, indicative of systemic bone formation, was lower in HFD-CR mice compared to controls in both male and female mice (Figure 5A). Caloric restriction reduced systemic bone resorption, evaluated with serum CTx, in females but not males (Figure 5B). Compared to control animals, caloric restriction decreased the number of osteoblasts per bone surface in trabecular bone in both male and female mice (Figure 5C). Osteoblast number was not changed by diet at the cortical surface. In both the cortical and trabecular compartments, the number of TRAP+ cells did not differ between diet groups; however, hematopoietic cells collected from male HFD-CR mice generated fewer osteoclasts *in vitro* compared to HFD animals (Figure S12). The size of cultured osteoclasts did not differ based on diet.

**Figure 5.**
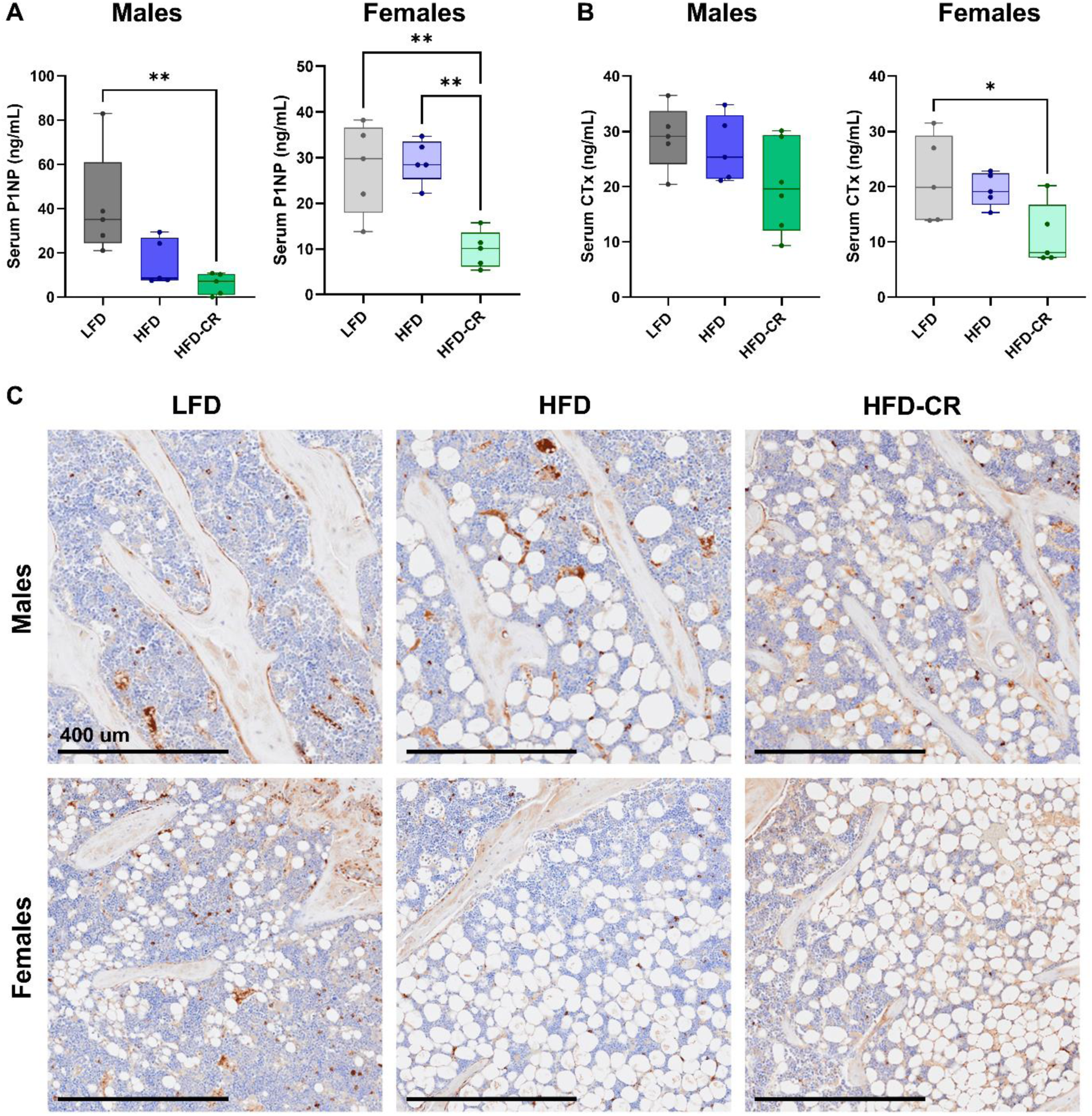
Calorie restriction reduced systemic bone formation and resorption. Serum collected at euthanasia was used to evaluate systemic bone formation and resorption with P1NP and CTx, respectively. (A) In male mice, HFD-CR reduced P1NP versus LFD. Compared to both LFD and HFD, caloric restriction reduced P1NP in female mice. (B) Diet did not change CTx in males. In females, HFD-CR reduced CTx compared to LFD mice. (C) Representative images of trabecular bone stained with procollagen-1 (brown). Scale bar = 400 um. Osteoblast number, normalized to bone surface, was reduced in HFD-CR compared to LFD mice. P1NP, CTx, and osteoblast number were evaluated by one-way ANOVA with factor of diet. Tukey post-hoc analysis determined differences between groups. p≤0.05 for all effects displayed. * indicates p≤0.05. ** indicates p≤0.01.

### Diet shifted the cortical metabolome in sex-specific manners

Both high-fat diet and calorie restriction altered the cortical bone metabolome. In both sexes, the metabolome of HFD mice was distinct from LFD (Figure S2). HFD-CR further shifted the metabolome away from LFD in both male and female mice. Specific pathways responsible for the diet-induced metabolomic differences were distinct between male and female mice. Following normalization to controls, unbiased hierarchical clustering of median metabolite intensity revealed distinct metabolomic differences between HFD and HFD-CR cortical bone (Figure 6). Pathways altered by diet generally mapped to amino acid or fatty acid metabolism.

**Figure 6.**
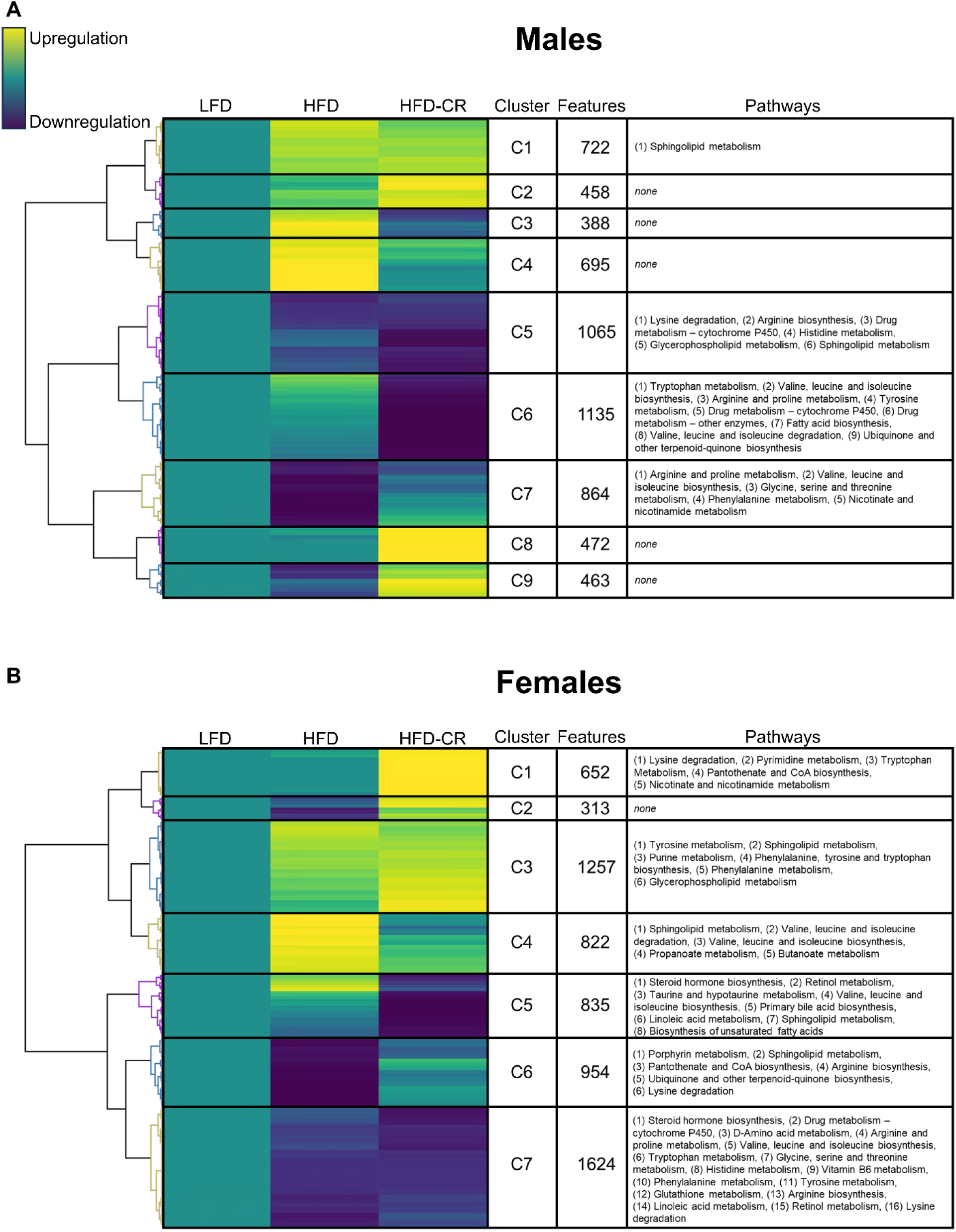
Cortical bone metabolome was altered by diet in a sex-dependent manner. Metabolites were extracted from cortical bone and untargeted metabolomics performed. Within each diet group, the median intensity of individual metabolites was calculated and normalized to the control group (LFD). Normalized intensities were hierarchically clustered. Each cluster was then correlated to pathways associated with individual metabolites. For both (A) male and (B) female mice, compared to LFD controls, yellow indicates upregulation of identified metabolites, teal indicates no difference from LFD, and blue indicates downregulation. The number of metabolites within each cluster was quantified (Features). Pathways related to clustered metabolites revealed that diet shifted the cortical bone metabolome in a sex-dependent manner.

In males, compared to LFD controls, most metabolic pathways identified were downregulated with HFD, HFD-CR, or in both groups. HFD reduced specific metabolic pathways that remained downregulated during subsequent caloric restriction; specifically, lysine degradation, arginine biosynthesis, arginine and proline metabolism, histidine metabolism, and glycerophospholipid metabolism, valine, leucine and isoleucine biosynthesis were downregulated in HFD and HFD-CR males compared to LFD controls. Compared to LFD, HFD males also had reduction of metabolic pathways that were generally not altered in HFD-CR mice, including glycine, serine, and threonine metabolism, phenylalanine metabolism, and nicotinate and nicotinamide metabolism. In males, HFD-CR, but not HFD, was associated with a downregulation of pathways related to tryptophan metabolism, tyrosine metabolism, ubiquinone and other terpenoid-quinone biosynthesis, valine, leucine and isoleucine degradation, and fatty acid biosynthesis.

In females, metabolic pathways were both up-and downregulated by dietary interventions. Many of the metabolic pathways shared by HFD and HFD-CR females were downregulated compared to LFD controls, including steroid hormone biosynthesis, D-Amino acid metabolism, arginine and proline metabolism, glycine serine and threonine metabolism, histidine metabolism, vitamin B6 metabolism, glutathione metabolism, arginine biosynthesis, linoleic acid metabolism, and retinol metabolism. The upregulation of some metabolic pathways by HFD were also upregulated in HFD-CR females, including purine metabolism, glycerophospholipid metabolism, and phenylalanine tyrosine and tryptophan biosynthesis. HFD-induced increases in propanoate metabolism, butanoate metabolism, and valine, leucine, and isoleucine degradation were not present in HFD-CR females. Similarly, HFD-induced decreases in porphyrin metabolism, and ubiquinone and other terpenoid-quinone biosynthesis were not recorded in HFD-CR females. Pantothenate and CoA biosynthesis were decreased in HFD mice but increased in HFD-CR females. Although not changed by HFD, HFD-CR females had increased pyrimidine metabolism and nicotinate and nicotinamide metabolism. Similarly, HFD-CR decreased taurine and hypotaurine metabolism, primary bile acid biosynthesis, and biosynthesis of fatty acids but these pathways were not changed by HFD.

## Discussion

Given the extent of the ongoing obesity epidemic, improved understanding of the effect of weight loss interventions on the skeleton are necessary. We demonstrated that in obese preclinical models, caloric restriction improved whole-body metabolism but further exacerbated bone loss. Reductions in bone formation and osteoblast number are likely responsible for diminished bone mass in calorie-restricted animals. In a sex-dependent manner, obesity and subsequent caloric restriction shifted the cortical bone metabolome, highlighting distinct changes to cellular metabolism within the skeleton in male and female mice.

Concurrent anabolic interventions may prevent bone loss during caloric restriction. In nonobese mice, simultaneous treatment with parathyroid hormone during caloric restriction attenuated trabecular bone loss^17^. Mechanical loading of the skeleton also can stimulate anabolic activity; in humans, resistance training exercise increased bone mineral density^32^. In our study, the increased ambulatory activity of the calorie-restricted mice did not prevent bone loss. Notably, increasing exercise through the addition of running wheels during the diet period also did not prevent calorie restriction-induced bone loss in nonobese mice^33^. In our study and others, during weight loss, ambulatory activity alone likely is not sufficient to produce enough anabolic stimuli for the skeleton; resistance training but not aerobic exercise attenuated bone loss in obese or overweight humans undergoing calorie restriction^34^.

Weight loss via caloric restriction versus the return to a normal diet elicits differential changes to the skeleton. In our study, caloric restriction negatively affected trabecular and cortical bone in both sexes whereas weight loss through the return to a normal diet altered bone morphology in a sex-dependent manner^9,22^. Previous work established that obese male mice returned to a normal chow diet had improved cancellous bone quality and similar cortical bone strength^9^. On the other hand, compared to lifelong obese mice, females returned to a normal diet retained obesity-induced trabecular deficits but had improved cortical bone strength^22^. After the return to a normal diet, the body mass of formerly obese mice was similar to controls^9,22^ but in our study, calorie restriction caused more weight loss, decreasing bodyweights below that of control groups. Differences in composition of control diets may also influence our ability to identify skeletal changes following weight loss; compared to standard chow diets, low fat diets are high in carbohydrate content and may exacerbate age-related bone loss^27^. Despite the potential for reduced bone mass in the control group, the use of sucrose-matched low fat diets are preferable when studying obesity^35^. Sucrose influences whole body metabolism and thus diets not matched in sucrose would introduce additional confounding variables^35^.

Different clinical strategies for weight loss may cause distinct changes to the skeleton. These interventions range from lifestyle changes to anti-obesity medications, and even surgical options. Although lifestyle interventions often have been the first-line clinical recommendations for weight loss, humans losing weight through diet and exercise typically regain weight after the intervention period^36^. The recent revolution in anti-obesity medications has expanded weight loss options for patients. Glucagon-like peptide-1 receptor agonists reduce appetite and food intake, thereby eliciting weight loss^37^. Early clinical data suggest that semaglutide causes cortical bone loss in humans^38^. For individuals who do not achieve weight loss with lifestyle interventions, anti-obesity medications may offer more skeletal protection compared to surgical interventions. Bariatric surgery has well established negative effects on bone in both humans^39,40^ and mouse models^41,42^, with preclinical studies suggesting that bone loss following bariatric surgery is greater than any potential bone loss due to anti-obesity medications^43^. Future work should evaluate potential differences in the skeleton following weight loss with caloric restriction versus therapeutic interventions.

The changes in metabolism of bone cells during obesity and subsequent weight loss likely drives the morphological changes recorded in our study, and highlights the importance of evaluating both male and female mice. The sex-specific responses of the cortical metabolome to dietary interventions reported here build on previous work, which established sexually dimorphic metabotypes in cortical bone^31^. In our study, within the cortical bone of male mice, caloric restriction after obesity reduced tryptophan metabolism, which has previously been linked to bone loss^44^. In females, the calorie restriction-induced downregulation of pathways related to taurine metabolism likely reflects altered osteocyte metabolism; taurine is a metabolite produced by osteocytes that positively regulates bone formation^45^. In both male and female mice, caloric restriction after obesity decreased fatty acid biosynthesis, which may have directly contributed to the reductions in osteoblast activity. Fatty acids are critical for osteoblast function, especially during nutrient limitation^46^. Notably, specific weight loss strategies likely elicit distinct changes to the metabolism within the bone; treatment with the SGLT2 inhibitor canagliflozin also causes weight loss, but within cortical bone, tryptophan catabolism was increased^47^.

Future work should address the limitations of our research. Individual bone cells rely on specific bioenergetic programs^48^. Although the cortical sample used for metabolomic analysis in this study was likely composed primarily of bone-resident osteocytes, we were unable to assign metabolites to specific cell types. Further, cortical and trabecular bone have distinct transcriptional signatures^49^, and thus may also rely on different bioenergetic programs. Although we identified calorie restriction-induced deficits in bone morphology and bone formation, we did directly evaluate bone strength. The translation of bone loss to increased fracture risk should be confirmed in future studies. Bone marrow adiposity has also been linked to overall skeletal health^50^. We measured increased bone marrow adiposity due to obesity and subsequent caloric restriction, but our small sample size may have precluded us from identifying any additive effects of weight loss after obesity.

This research highlights the negative effects of obesity and subsequent caloric restriction on the skeleton. We determined that in both male and female mice, caloric restriction reduced cortical bone mass and trabecular thickness. Notably, the cortical bone metabolome was changed in obese mice compared to control animals, but further shifted by subsequent caloric restriction. Diet-induced metabolic shifts were sex-specific, highlighting the potential for distinct mechanisms to contribute to calorie restriction-induced bone loss in male versus female mice. Importantly, this work highlights potential clinical concerns surrounding weight loss after obesity; although calorie restriction may improve whole body metabolic outcomes, fracture risk will likely increase.

## Ethics Approval/Institutional Review Board Statement

The animal study protocol was approved by the Institutional Animal Care and Use Committee (IACUC) of MaineHealth Institute for Research (protocol #2209).

## Acknowledgements

We thank Peter Caradonna for assistance with histology and Jen Daruszka for assistance with animal work.

