## Supplemental Material for "In C57Bl6 Mice, Obesity and Subsequent Weight Loss Negatively Affected the Skeleton and Shifted the Cortical Bone Metabolome"

**Supplemental Materials**

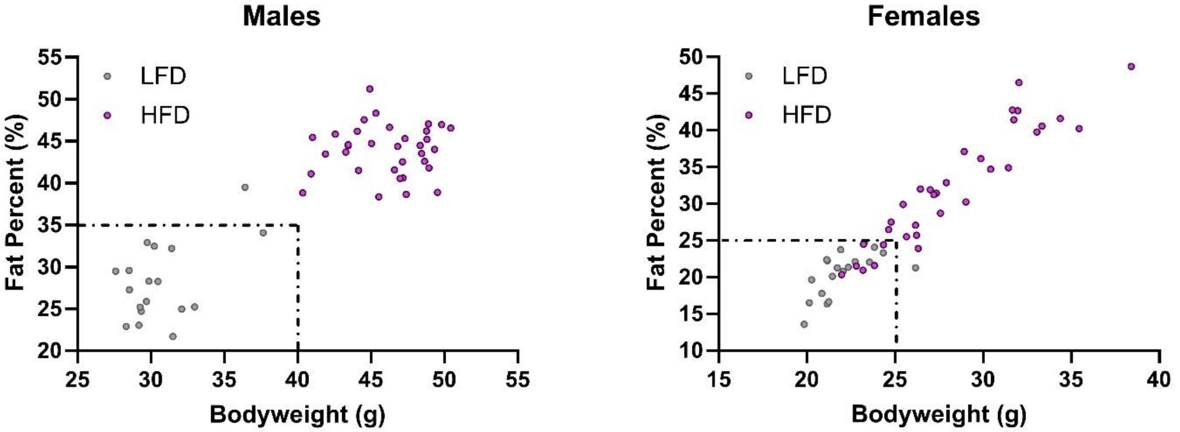


**Figure S1. Bodyweight and fat mass were used to identify non-obese mice following the High Fat Feeding Phase.** (A) In males, obesity was defined as animals weighing at least 40 g or having greater than 35% areal fat mass. Fat mass was determined via DXA scans. All male mice consuming high fat diet met obesity criteria. (B) In females, obesity was defined as animals weighing at least 25 g or having greater than 25% areal fat mass. Of the 34 females consuming high fat diet, 7 were not considered to be obese at the conclusion of the High Fat Feeding Phase and were removed from the study.

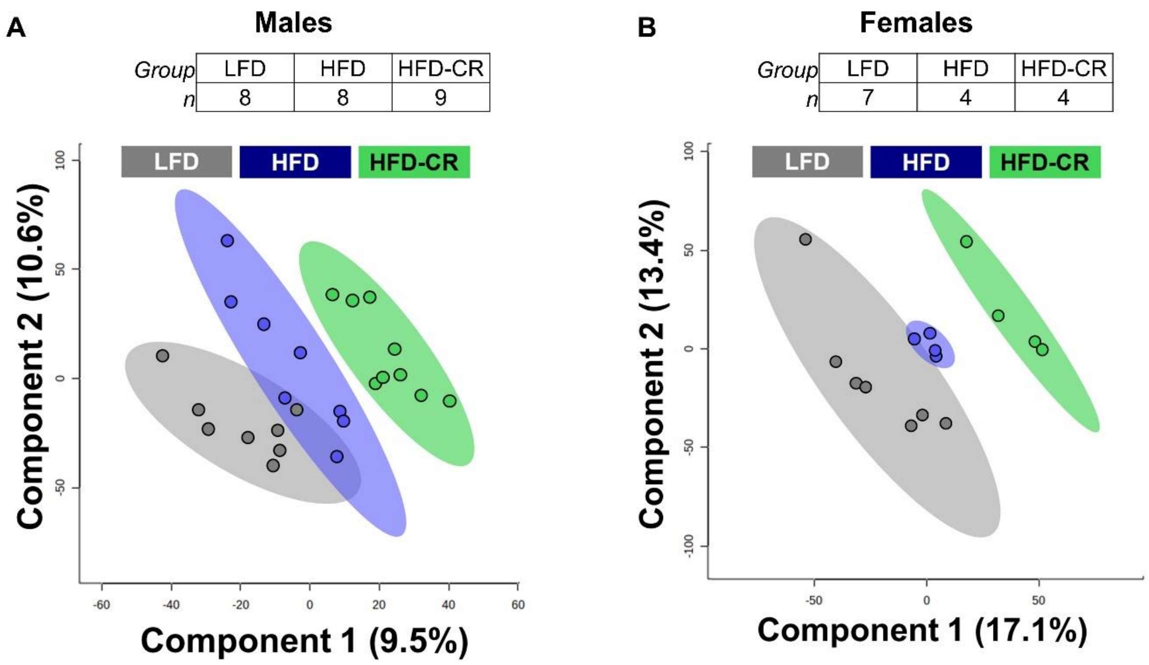


**Figure S2. Diet shifted cortical bone metabolome in both male and female mice.** Metabolites from cortical bone were extracted and identified with unbiased LC-MS. Using partial least-squares discriminant analysis, dimensionality was reduced, revealing that in both (A) males and (B) females, the cortical bone metabolome of HFD mice was dissimilar from LFD, and subsequent caloric restriction even further altered the cortical bone metabotype. Analysis performed in Metaboanalyst. n=4-9 per group.

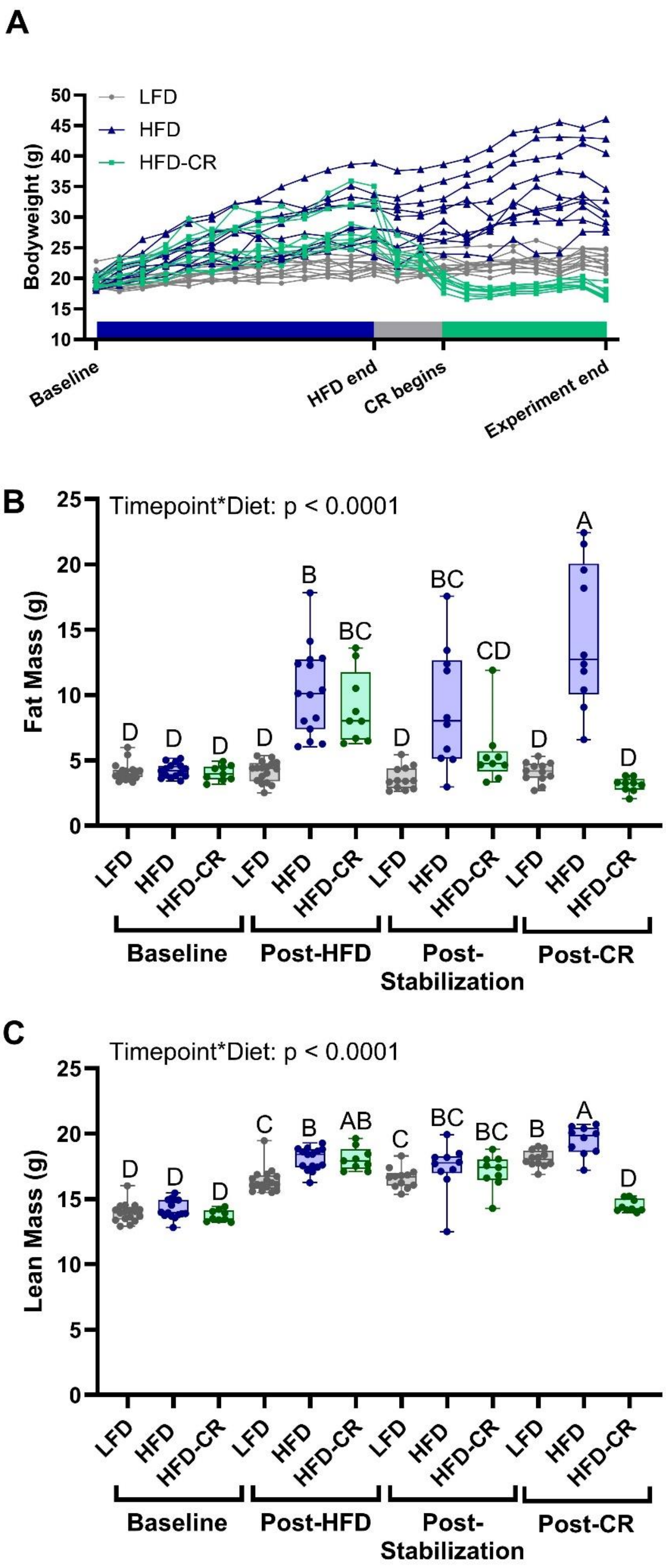
A
LFD
HFD
HFD-CR
Bodyweight (g)
50
45
40
35
30
25
20
15
10
Baseline
HFD end
CR begins
Experiment end
B
Timepoint*Diet: p < 0.0001
Fat Mass (g)
25
20
15
10
5
0
D D D D B BC D BC CD D A D
LFD HFD HFD-CR LFD HFD HFD-CR LFD HFD HFD-CR LFD HFD HFD-CR
Baseline
Post-HFD
Post-Stabilization
Post-CR
C
Timepoint*Diet: p < 0.0001
Lean Mass (g)
25
20
15
10
5
0
D D D C B AB C BC BC B A D
LFD HFD HFD-CR LFD HFD HFD-CR LFD HFD HFD-CR LFD HFD HFD-CR
Baseline
Post-HFD
Post-Stabilization
Post-CR

**Figure S3. In female mice, body mass increased during high fat feeding and decreased during caloric restriction.** (A) During the High Fat Feeding Phase, females gained bodyweight. During the Caloric Restriction Phase, calorie restricted mice lost weight. (B) High Fat Feeding increased fat mass, evaluated by DXA. Subsequent caloric restriction decreased fat mass. (C) Lean mass was increased by HFD but decreased following caloric restriction. Data displayed includes females that were not excluded prior to the end of the study (see details in Methods). Fat and lean mass were analyzed by two-way ANOVA with factors of timepoint and diet. Significant terms are displayed. Tukey post-hoc analysis determined differences between groups. $p \leq 0.05$ for all effects displayed. Letters denote statistical differences: A>B>C>D.

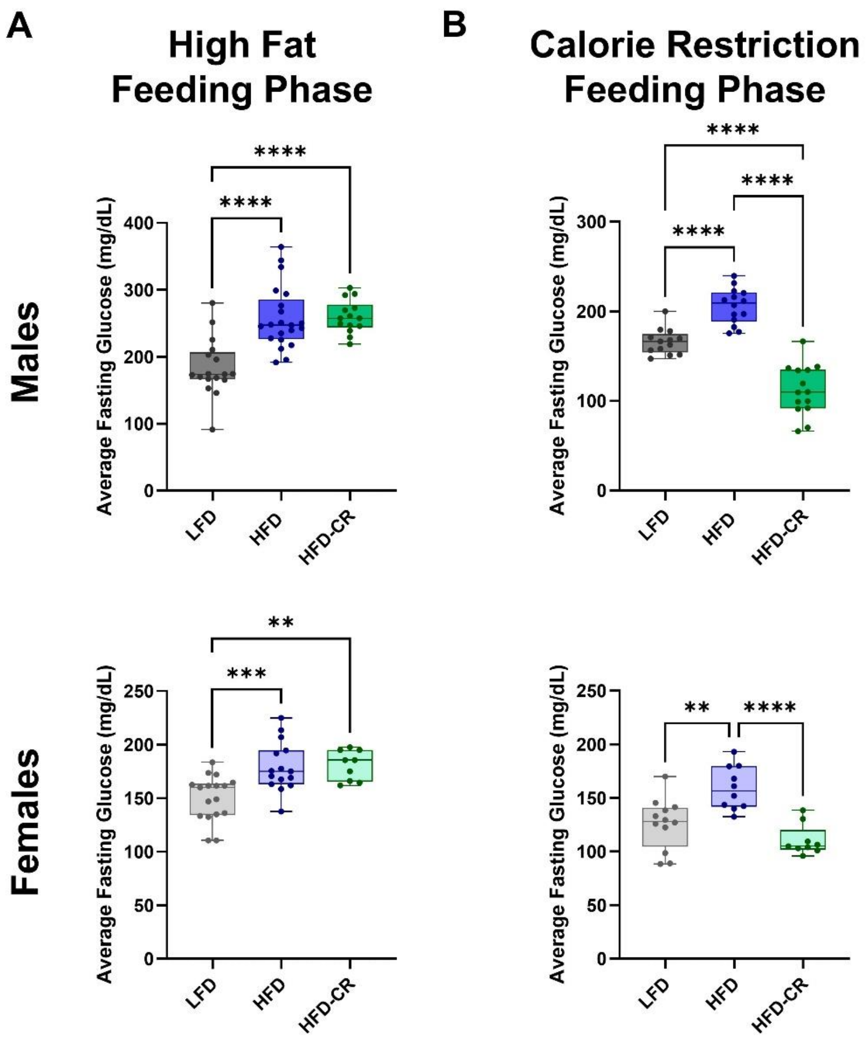


**Figure S4. Fasting blood glucose was increased by high fat feeding, but decreased by subsequent caloric restriction.** Average fasting blood glucose was calculated using baseline readings for glucose and insulin tolerance tests. Animals were fasted for 6 hours prior to measurements. (A) At the conclusion of the High Fat Feeding Phase, groups receiving HFD had increased fasting blood glucose compared to LFD controls. (B) Subsequent caloric restriction decreased fasting blood glucose in HFD-CR compared to HFD mice. Average fasting blood glucose was evaluated by one-way ANOVA with factor of diet. Tukey post-hoc analysis determined differences between groups. $p \leq 0.05$ for all effects displayed. ** indicates $p \leq 0.01$. **** indicates $p \leq 0.0001$.

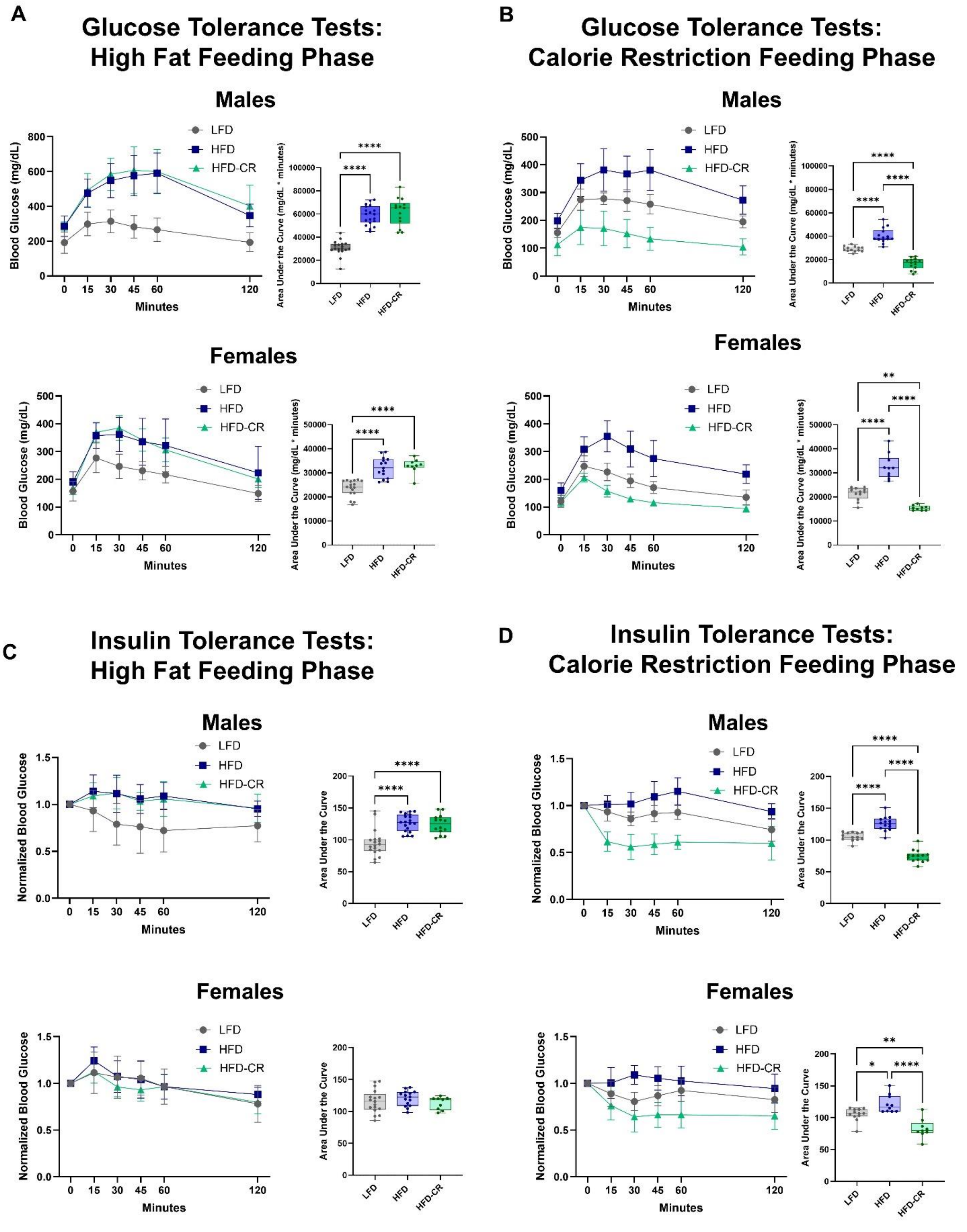
A
Glucose Tolerance Tests:
High Fat Feeding Phase
Males
Females
B
Glucose Tolerance Tests:
Calorie Restriction Feeding Phase
Males
Females
C
Insulin Tolerance Tests:
High Fat Feeding Phase
Males
Females
D
Insulin Tolerance Tests:
Calorie Restriction Feeding Phase
Males
Females
LFD
HFD
HFD-CR
Blood Glucose (mg/dL)
Normalized Blood Glucose
Minutes
Area Under the Curve (mg/dL * minutes)
Area Under the Curve

**Figure S5. Glucose and insulin tolerance were worsened by high fat feeding, but improved by subsequent caloric restriction.** Average fasting blood glucose was calculated using baseline readings for glucose and insulin tolerance tests. Animals were fasted for 6 hours prior to measurements. (A) At the conclusion of the High Fat Feeding Phase, groups receiving HFD had worsened glucose tolerance compared to LFD controls. (B) Subsequent caloric restriction improved glucose tolerance in HFD-CR compared to both HFD and LFD mice. (C) At the conclusion of the High Fat Feeding Phase, males receiving HFD had worsened insulin tolerance compared to LFD controls. In females, insulin tolerance did not differ between diet groups at the end of the High Fat Feeding Phase. (D) In male mice, subsequent caloric restriction improved insulin tolerance in HFD-CR compared to both HFD and LFD animals. At the conclusion of the Caloric Restriction Phase, compared to LFD mice, insulin tolerance was worsened in HFD females but improved in HFD-CR. Line graphs display mean and standard deviation for each group. Area under the curve was calculated individually for each mouse and evaluated by one-way ANOVA with factor of diet. Tukey post-hoc analysis determined differences between groups. $p \leq 0.05$ for all effects displayed. * indicates $p \leq 0.05$. ** indicates $p \leq 0.01$. **** indicates $p \leq 0.0001$.

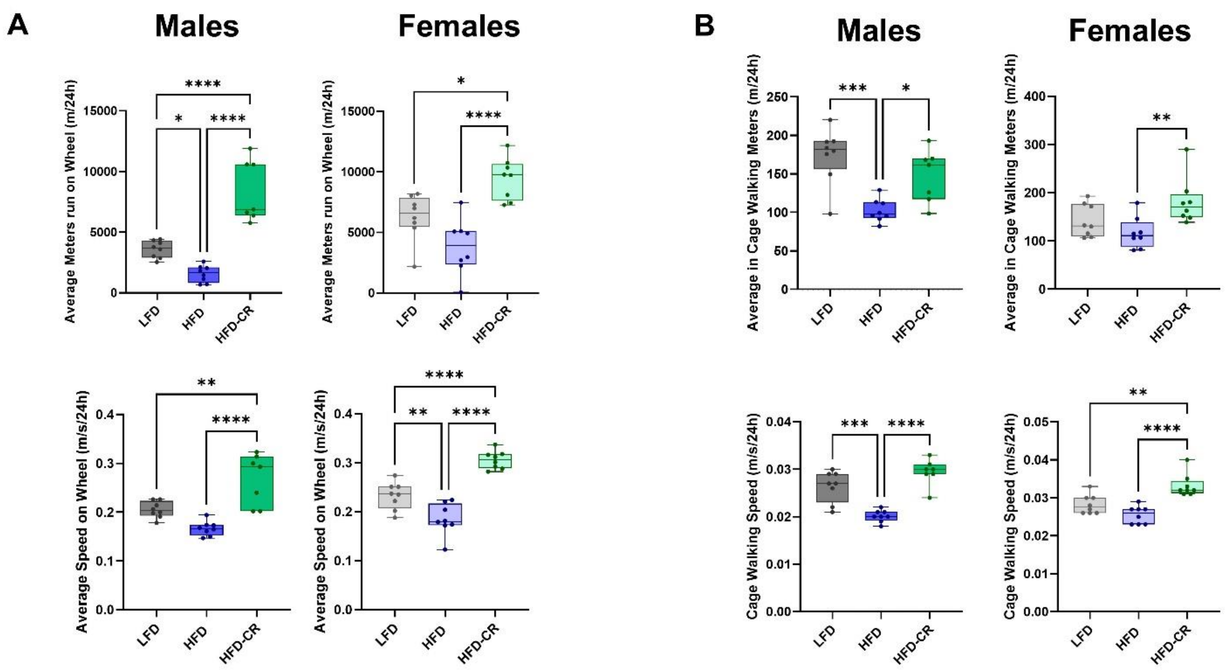


**Figure S6. Calorie restriction altered ambulatory patterns in both male and female mice.** Ambulatory activity was quantified in metabolic cages at the end of the Caloric Restriction Phase. (A) In both sexes, HFD-CR animals ran further and faster on the wheel compared to both LFD and HFD mice. (B) Compared to HFD, HFD-CR mice walked further and faster within the cage. Ambulatory activity was evaluated by one-way ANOVA with factor of diet. Tukey post-hoc analysis determined differences between groups. $p \leq 0.05$ for all effects displayed. * indicates $p \leq 0.05$. ** indicates $p \leq 0.01$. *** indicates $p \leq 0.001$. **** indicates $p \leq 0.0001$.

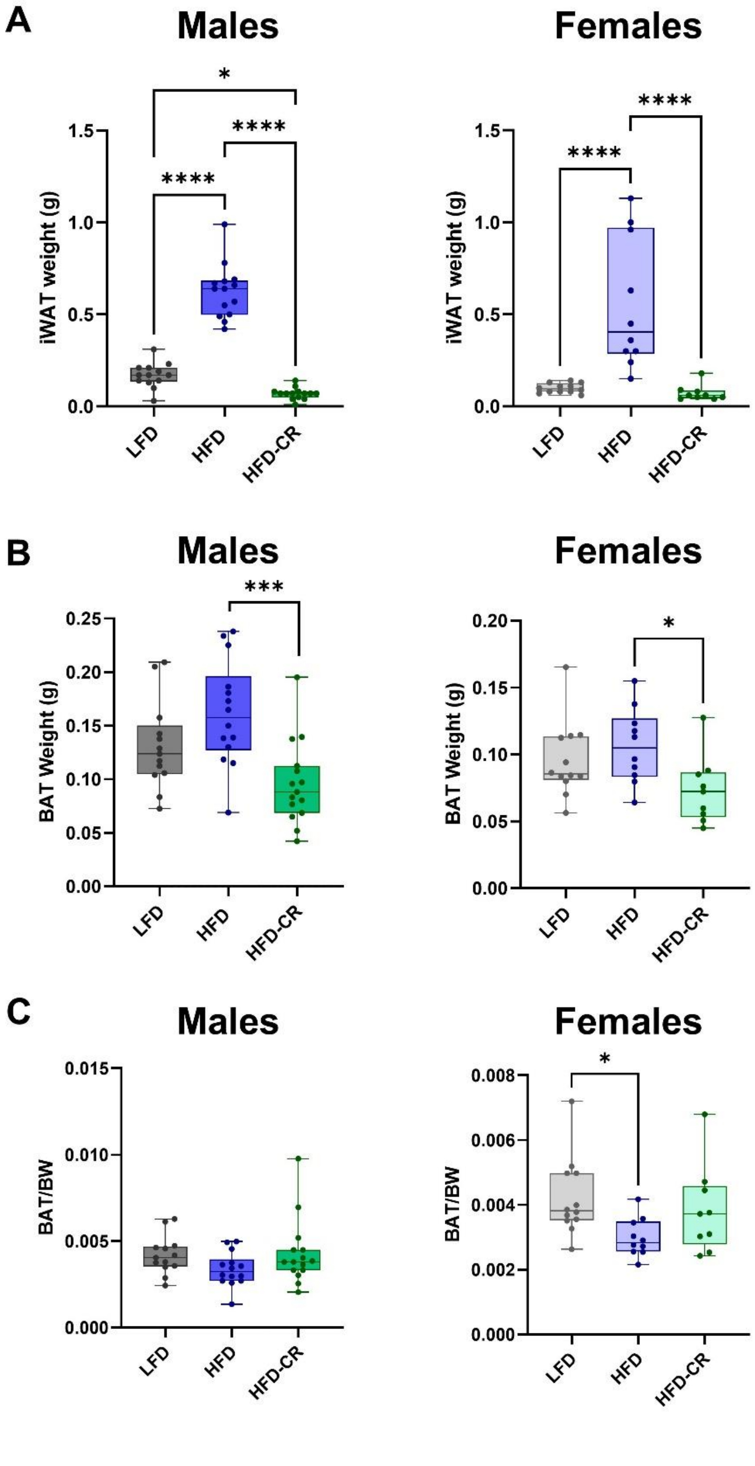


**Figure S7. Inguinal adipose depot weights were increased in HFD mice but decreased by subsequent caloric restriction.** Inguinal and brown adipose depots were collected and weighed at euthanasia. (A) In both sexes, HFD increased inguinal white adipose tissue (iWAT) weight compared to

LFD mice. HFD-CR animals had smaller iWAT versus HFD mice. (B) In males, HFD-CR decreased brown adipose tissue (BAT) weight. BAT weight was not changed by diet in females. (C) When normalized to bodyweight, BAT weights were not changed by diet in either sex. Adipose depot weights were evaluated by one-way ANOVA with factor of diet. Tukey post-hoc analysis determined differences between groups. p≤0.05 for all effects displayed. * indicates p≤0.05. *** indicates p≤0.001. **** indicates p≤0.0001.

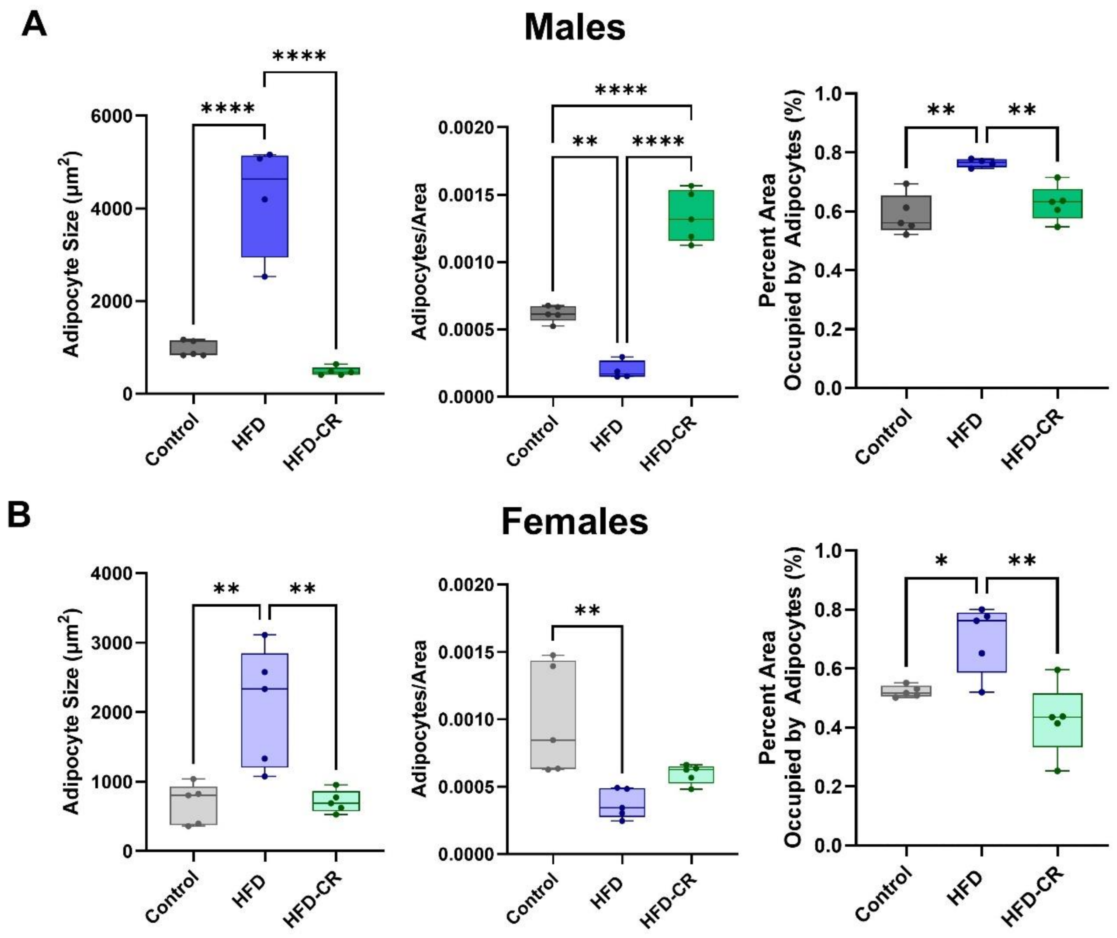


**Figure S8. Inguinal adipocyte size was increased by HFD and decreased by subsequent caloric restriction.** Adiposity was quantified by hematoxylin and eosin-stained sections. Adipocyte size and number were quantified in a representative region of interest in both (A) males and (B) females. In both sexes, adipocyte size was increased by HFD. HFD reduced the number of adipocytes within the area analyzed but increased the percent area occupied by adipocytes. Adiposity measures were evaluated by one-way ANOVA with factor of diet. Tukey post-hoc analysis determined differences between groups. $p \leq 0.05$ for all effects displayed. * indicates $p \leq 0.05$. ** indicates $p \leq 0.01$. **** indicates $p \leq 0.0001$.

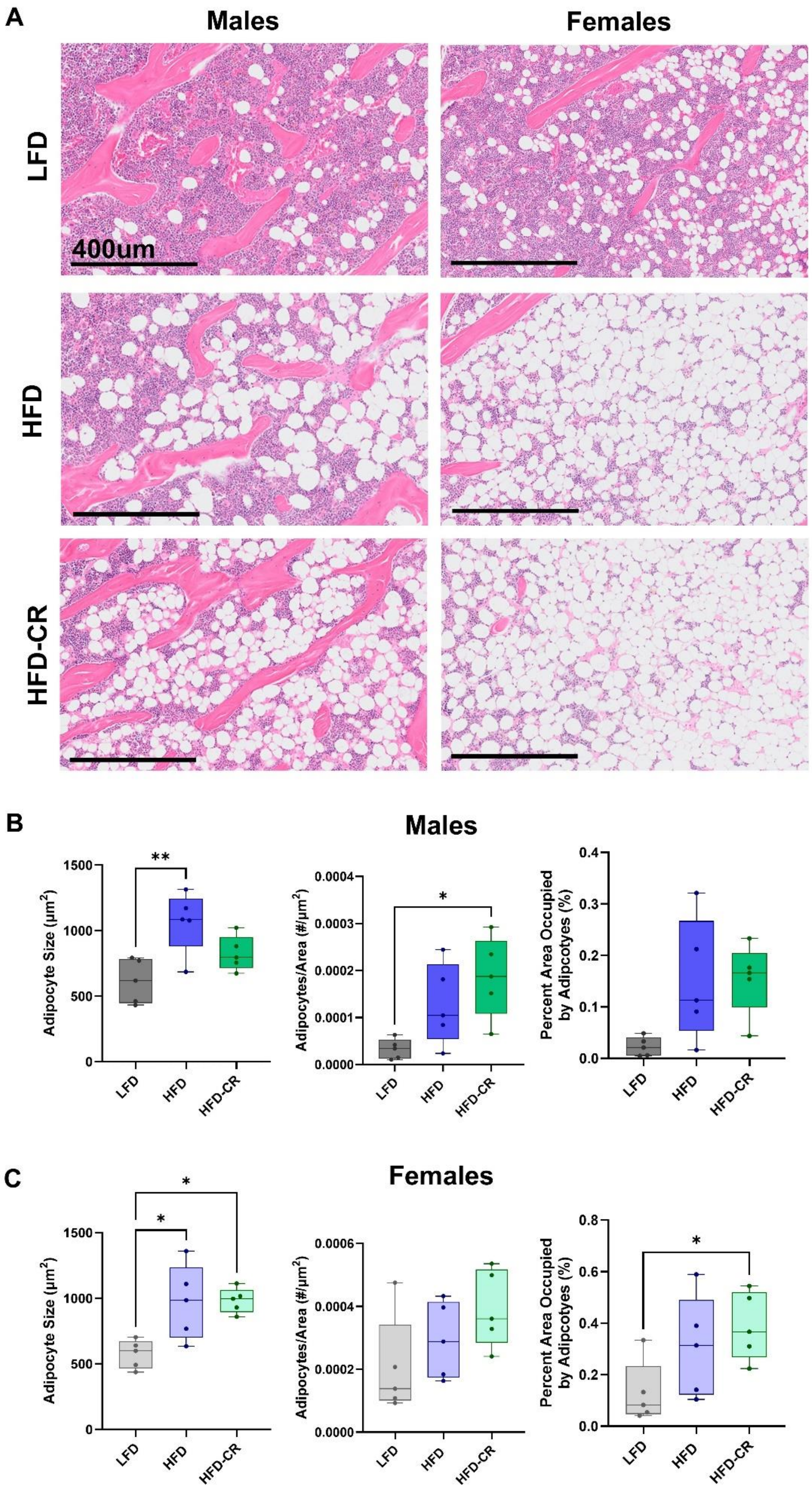
A
Males
Females
LFD
HFD
HFD-CR
400um
B
Males
Adipocyte Size (µm²)
Adipocytes/Area (#/µm²)
Percent Area Occupied by Adipcotyes (%)
LFD
HFD
HFD-CR
C
Females
Adipocyte Size (µm²)
Adipocytes/Area (#/µm²)
Percent Area Occupied by Adipcotyes (%)
LFD
HFD
HFD-CR

**Figure S9. Bone marrow adipocyte size was increased by HFD but not changed by subsequent caloric restriction.** Adiposity was quantified by hematoxylin and eosin-stained sections. (A) Representative images. Scale bar = 400 um. (B) In male mice, HFD increased bone marrow adipocyte size compared to LFD animals. HFD-CR males had more bone marrow adipocytes than LFD. The percent area occupied by bone marrow adipocytes was not altered by diet in male mice. (C) Both HFD and HFD-CR females had larger bone marrow adipocytes compared to LFD. The number of bone marrow adipocytes was not altered by diet in female mice. Compared to LFD, HFD-CR increased the percent area occupied by bone marrow adipocytes. Adiposity measures were evaluated by one-way ANOVA with factor of diet. Tukey post-hoc analysis determined differences between groups. $p \leq 0.05$ for all effects displayed. * indicates $p \leq 0.05$. ** indicates $p \leq 0.01$.

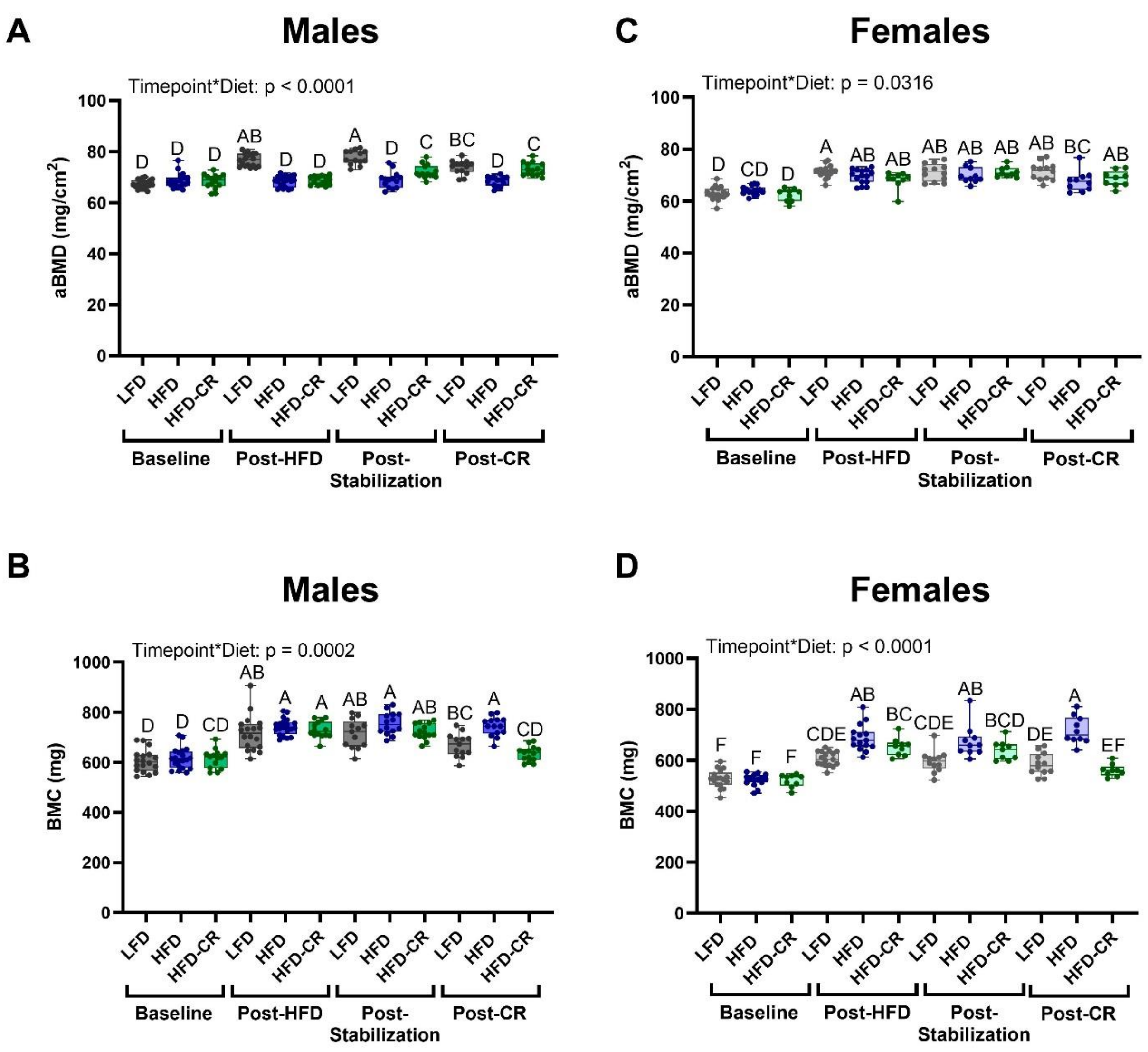


**Figure S10. Obesity and subsequent caloric restriction affected areal bone mineral density and bone mineral content.** Areal bone mineral density (aBMD) and bone mineral content (BMC) was quantified at baseline and at the end of each diet phase. (A) At the end of the High Fat Feeding Phase, males receiving a high fat diet had lower aBMD compared to LFD. After the Stabilization Phase, HFD-CR mice had increased aBMD compared to HFD. aBMD remained elevated in HFD-CR males compared to HFD at the end of the Caloric Restriction Phase. (B) In male mice, BMC did not differ between diet groups until the end of the Caloric Restriction Phase; HFD males had greater BMC compared to LFD and HFD-CR. (C) In females, aBMD was not statistically different between groups at any phase of the diet. aBMD increased over time in all diet groups throughout the study. (D) At the conclusion of the study, HFD females had greater BMC compared to LFD and HFD-CR mice. aBMD and BMC were analyzed by two-way ANOVA with factors of timepoint and diet. Significant terms are displayed. Tukey post-hoc analysis determined differences between groups. $p \leq 0.05$ for all effects displayed. Letters denote statistical differences: A>B>C>D>E>F.

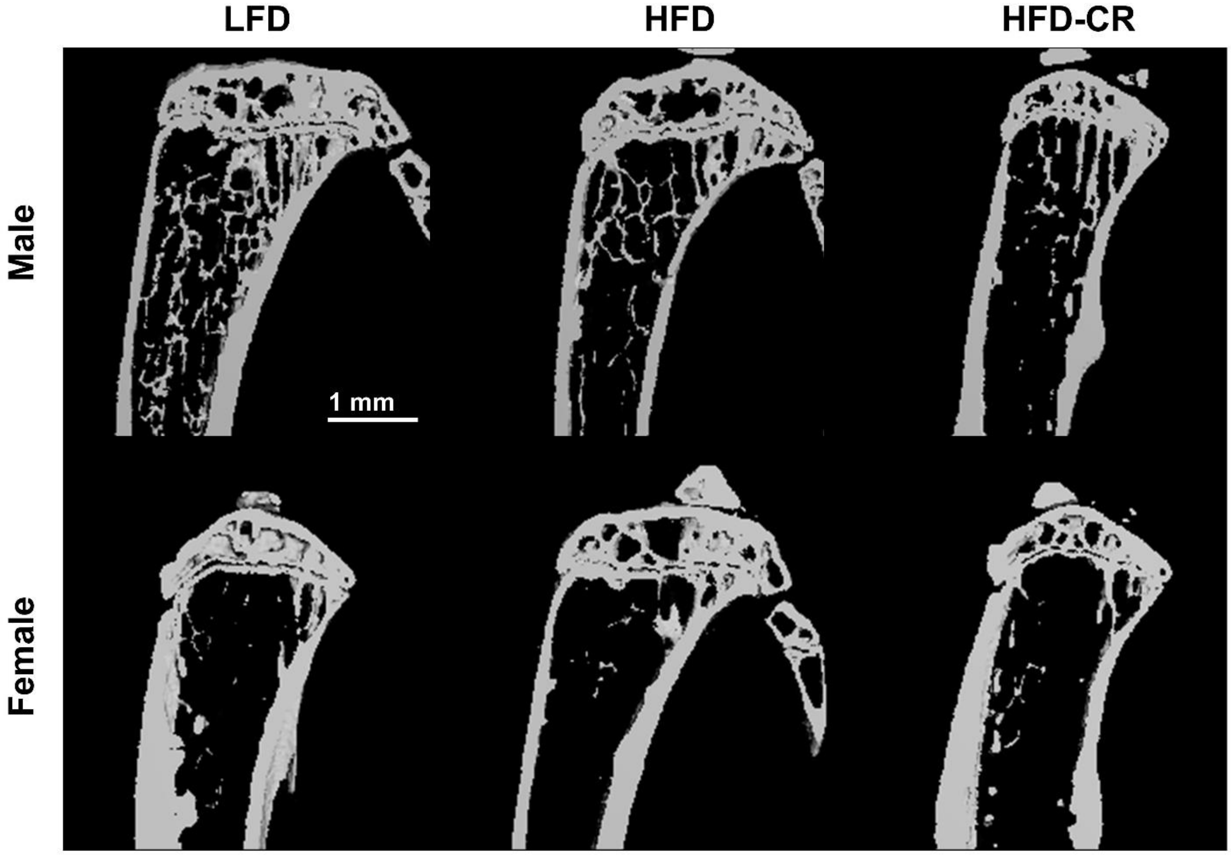


**Figure S11. Diet altered bone morphology in both male and female mice.** Representative µCT images. Scale bar = 1 mm.

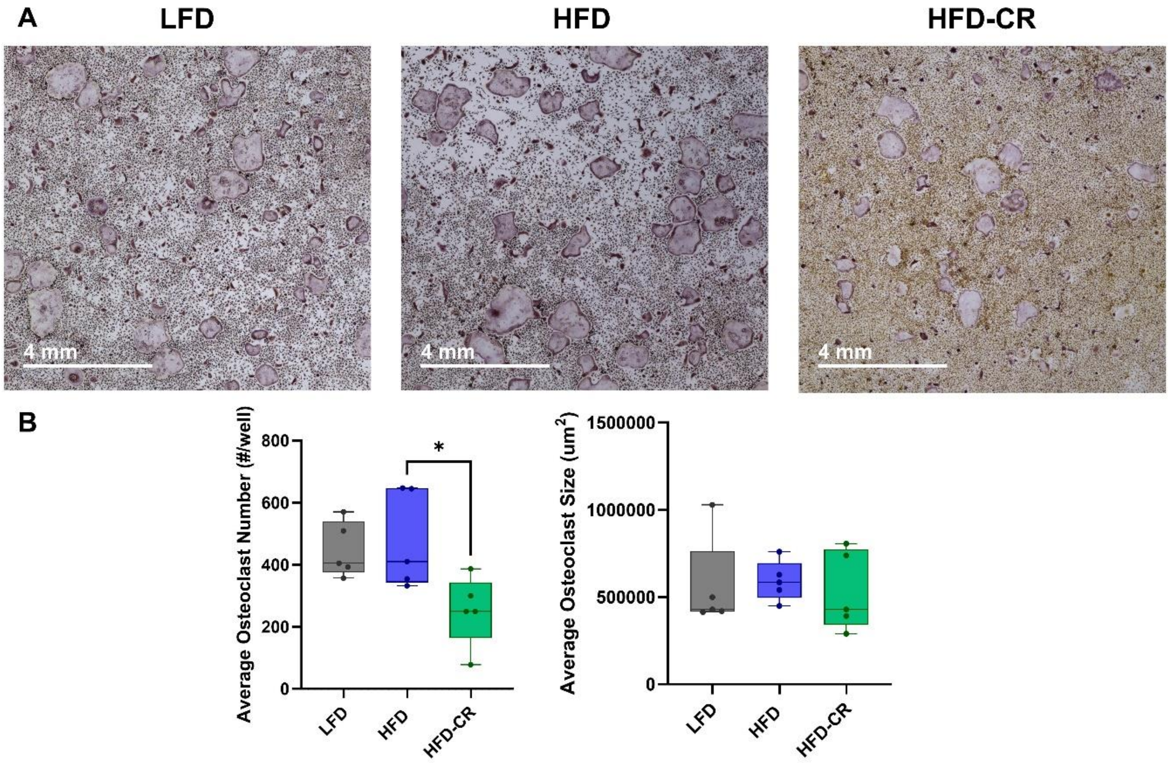


**Figure S12. Caloric restriction negatively affected osteoclastogenesis in male mice.** At the conclusion of the study, hematopoietic stem cells were collected from the bone marrow of long bones and cultured in osteoclast differentiation media. Five animals per group were included for analysis and cultured individually. Following 7 days in culture, cells were stained for TRAP and imaged. (A) Representative images. Scale bar = 4 mm. (B) Using two wells per mouse, osteoclast number and size were quantified in ImageJ. Osteoclasts were defined as TRAP+ cells containing 3 or more nuclei. Compared to HFD, caloric restriction reduced the number of differentiated osteoclasts, but did not change osteoclast size. Osteoclast number and size were evaluated by one-way ANOVA with factor of diet. Tukey post-hoc analysis determined differences between groups. $p \leq 0.05$ for all effects displayed. * indicates $p \leq 0.05$.

**Table S1. At the end of the High Fat Feeding Phase, adiposity was increased in mice receiving high fat diet.** At the conclusion of the High Fat Feeding Phase, a subset of mice was euthanized to confirm obesity phenotype (n=5/group). Adipose depots were collected and weighed. Adiposity of inguinal and bone marrow adipose tissues were quantified in ImageJ. Average and standard deviation are reported for each measure. Differences were determined using unpaired t-tests. Significant p-values are bolded and indicated by *.

| | Males | | | Females | | |
|---|---|---|---|---|---|---|
| *Parameter* | Control | HFD | p-value | Control | HFD | p-value |
| iWAT weight (g) | 0.146 ± 0.013 | 0.578 ± 0.152 | **0.0002*** | 0.088 ± 0.040 | 0.266 ± 0.061 | **0.0006*** |
| BAT weight (g) | 0.076 ± 0.011 | 0.096± 0.030 | 0.2213 | 0.050 ± 0.016 | 0.071 ± 0.026 | 0.1645 |
| BAT/BW | 0.0025 ± 0.0003 | 0.0022 ± 0.0007 | 0.3779 | 0.0022 ± 0.0007 | 0.0023 ± 0.0008 | 0.8294 |
| BMAT Adipocyte Size (μm$^2$) | 546.62 ± 165.52 | 965.44 ± 76.63 | **0.0026*** | 523.95 ± 90.66 | 853.29 ± 103.71 | **.0015*** |
| BMAT Adipocytes/Area (#/μm$^2$) | 0.000064 ± 0.000050 | 0.000104 ± 0.000046 | 0.2259 | 0.000097 ± 0.000025 | 0.000207 ± 0.000110 | 0.0882 |
| BMAT Percent Area Occupied by Adipocytes (%) | 0.041 ± 0.040 | 0.101 ± 0.045 | 0.0598 | 0.053 ± 0.022 | 0.185 ± 0.113 | 0.0582 |
| iWAT Adipocyte Size (μm$^2$) | 1267.83 ± 473.70 | 4094.14 ± 1962.51 | **.0303*** | 920.71 ± 229.37 | 2019.18 ± 314.48 | **.0003*** |
| iWAT Adipocytes/Area (#/μm$^2$) | 0.000548 ± 0.000121 | 0.000199 ± 0.000073 | **0.0011*** | 0.000730 ± 0.000244 | 0.000394 ± 0.000051 | **.0354*** |
| iWAT Percent Area Occupied by Adipocytes (%) | 0.646 ± 0.103 | 0.687 ± 0.161 | 0.6415 | 06295 ± 0.057 | 0.767 ± 0.023 | **.0034*** |

**Table S2. The skeleton was negatively affected by high fat diet at the conclusion of the High Fat Feeding Phase.** At the conclusion of the High Fat Feeding Phase, a subset of mice was euthanized to confirm obesity phenotype (n=5/group). Long bones were collected and μCT and dynamic histomorphometry performed. Average and standard deviation are reported for each measure. Differences were determined using unpaired t-tests. Significant p-values are bolded and indicated by *.

| | Males | | | Females | | |
|---|---|---|---|---|---|---|
| *Parameter* | Control | HFD | p-value | Control | HFD | p-value |
| Cortical Area (mm$^2$) | 0.71 ± 0.06 | 0.65 ± 0.05 | 0.1361 | 0.57 ± 0.01 | 0.60 ± 0.01 | **0.0134*** |
| Cortical Thickness (mm) | 0.22 ± 0.01 | 0.20 ± 0.01 | **0.0194*** | 0.19 ± 0.01 | 0.19 ± 0.01 | 0.4783 |
| Cortical Area/Total Area (%) | 60.04 ± 1.01 | 58.68 ± 1.94 | 0.2111 | 58.39 ± 1.67 | 58.28 ± 1.16 | 0.9033 |
| Marrow Area (mm$^2$) | 0.47 ± 0.04 | 0.46 ± 0.06 | 0.7001 | 0.41 ± 0.03 | 0.43 ± 0.02 | 0.3086 |
| Maximum Moment of Inertia (mm$^4$) | 0.13 ± 0.02 | 0.11 ± 0.02 | 0.3163 | 0.08 ± 0.01 | 0.08 ± 0 | 0.1382 |
| Cortical TMD (mgHA/cm$^3$) | 1267.27 ± 12.35 | 1273.54 ± 13.49 | 0.4650 | 1283.02 ± 11.75 | 1275.46 ± 5.18 | 0.2398 |
| Tibial Length (mm) | 18.30 ± 0.19 | 18.11 ± 0.15 | 0.1179 | 17.82 ± 0.19 | 17.97 ± 0.19 | 0.2408 |
| Periosteal BFR (um/day) | 0.21 ± 0.22 | 0.35 ± 0.07 | 0.2562 | 0.22 ± 0.04 | 0.08 ± 0.12 | 0.0570 |
| Periosteal MS/BS (%) | 33.25 ± 16.30 | 35.53 ± 6.09 | 0.7812 | 26.97 ± 3.28 | 18.02 ± 12.11 | 0.1769 |
| Periosteal MAR (um/day) | 0.52 ± 0.47 | 0.98 ± 0.17 | 0.0936 | 0.81 ± 0.16 | 0.31 ± 0.43 | 0.0565 |
| Endosteal BFR (um/day) | 0.53 ± 0.18 | 0.66 ± 0.22 | 0.3570 | 0.77 ± 0.19 | 0.78 ± 0.26 | 0.9802 |
| Endosteal MS/BS (%) | 49.87 ± 6.52 | 51.00 ± 15.40 | 0.8859 | 78.44 ± 7.70 | 71.89 ± 6.86 | 0.1935 |
| Endosteal MAR (um/day) | 1.04 ± 0.27 | 1.29 ± 0.21 | 0.1485 | 0.97 ± 0.17 | 1.07 ± 0.27 | 0.5401 |
| BV/TV (%) | 21.07 ± 6.00 | 15.60 ± 3.38 | 0.1236 | 10.00 ± 0.73 | 10.15 ± 1.05 | 0.8036 |
| Trabecular Thickness (mm) | 0.06 ± 0 | 0.05 ± 0 | **0.0449*** | 0.06 ± 0 | 0.06 ± 0 | 0.7877 |
| Trabecular Sp. (mm) | 0.17 ± 0.02 | 0.18 ± 0.02 | 0.2594 | 0.29 ± 0.03 | 0.30 ± 0.02 | 0.7126 |

| | | | | | | |
|---|---|---|---|---|---|---|
| Trabecular Number (1/mm) | 5.68 ± 0.77 | 5.29 ± 0.49 | 0.3713 | 3.48 ± 0.28 | 3.40 ± 0.30 | 0.6368 |
| Trabecular TMD (mgHA/cm$^3$) | 215.44 ± 43.99 | 173.19 ± 25.58 | 0.1095 | 133.32 ± 6.39 | 131.39 ± 10.90 | 0.7432 |
| Trabecular BFR (um/day) | 0.15 ± 0.06 | 0.09 ± 0.02 | 0.0609 | 0.21 ± 0.03 | 0.15 ± 0.10 | 0.2679 |
| Trabecular MS/BS (%) | 21.83 ± 4.98 | 14.99 ± 4.71 | 0.0562 | 23.54 ± 3.15 | 20.25 ± 9.72 | 0.5046 |
| Trabecular MAR (um/day) | 0.68 ± 0.16 | 0.58 ± 0.08 | 0.2403 | 0.88 ± 0.11 | 0.72 ± 0.18 | 0.1288 |
